# A novel oxidase from *Alcaligenes* sp. HO-1 oxidizes hydroxylamine to N_2_

**DOI:** 10.1101/2020.08.20.256677

**Authors:** Meng-Ru Wu, Li-Li Miao, Ying Liu, Ting-Ting Hou, Guo-Min Ai, Lan Ma, Hai-Zhen Zhu, Ya-Xin Zhu, Xi-Yan Gao, Xin-Xin Qian, Ya-Ling Qin, Tong Wu, Xi-Hui Shen, Cheng-Ying Jiang, Craig W. Herbold, Michael Wagner, De-Feng Li, Zhi-Pei Liu, Shuang-Jiang Liu

**Affiliations:** State Key Laboratory of Microbial Resources, Institute of Microbiology, Chinese Academy of Sciences, Beijing, China; University of Chinese Academy of Sciences, Beijing, China; University of Vienna, Centre for Microbiology and Environmental Systems Science, Division of Microbial Ecology Vienna, Austria; Shaanxi Key Laboratory of Agricultural and Environmental Microbiology, College of Life Sciences, Northwest A&F University, Yangling, Shaanxi, People’s Republic of China; Center for Microbial Communities, Aalborg University, Aalborg, Denmark; Environmental Microbiology Research Center, Institute of Microbiology, Chinese Academy of Sciences, Beijing, China; Joint-Lab of Environmental Biotechnology, IMCAS-RECEES, Beijing, China

**Author notes:** Correspondence to: Shuang-Jiang Liu, Zhi-Pei Liu, or De-Feng Li. These authors contributed equally to this work (Meng-Ru Wu, Li-Li Miao, Ying Liu, Ting-Ting Hou).

## Abstract

Hydroxylamine is a key intermediate of microbial ammonia oxidation and plays an important role in the biogeochemical cycling of N-compounds. Hydroxylamine is oxidized to NO or N_2_O by hydroxylamine oxidases or cytochrome P460 from heterotrophic or autotrophic bacteria, but its enzymatic oxidation to N_2_ has not yet been observed. Here, we report on the discovery of a novel oxidase that converts hydroxylamine to N_2_ from the newly isolated heterotrophic nitrifier *Alcaligenes* strain HO-1. Strain HO-1 accumulated hydroxylamine and produced N_2_ from ammonia oxidation. Using transcriptome analysis and heterologous expression via fosmid library screening, we identified three genes (*dnfABC*) of strain HO-1 that enabled *E. coli* cells not only to produce hydroxylamine from ^15^N-labelled ammonium but also to further convert it to ^15^N_2_. The three genes were individually cloned and expressed, and their translational products DnfA, DnfB, and DnfC were purified. *In vitro* DnfA bound to hydroxylamine and catalyzed the conversion of hydroxylamine to N_2_ in the presence of FAD, NADH and O_2_. Thus, DnfA was identified as a novel hydroxylamine oxidase and catalyzed a previously unknown N-N bond forming reaction with a yet-to-be discovered mechanism. DnfA homologs were detected in different bacterial groups, suggesting that hydroxylamine oxidation to nitrogen might occur in additional microbial taxa.

## Introduction

Hydroxylamine is formed by diverse microorganisms, including ammonia-oxidizing bacteria (AOB) and archaea (AOA) (1, 2), complete ammonia oxidizers (3, 4), anaerobic ammonia oxidizers (5), heterotrophic nitrifiers (6), fungi (7), and methane-oxidizing bacteria (8). Hydroxylamine is not only an important intermediate/side product in various microbial physiological pathways, but its production also has significant implications for the health of the environment. Both biological and non-biological processes are involved in the conversion of hydroxylamine to N_2_O (4, 9-11), which is an important greenhouse gas. Hydroxylamine production by soil microbes contributes to the formation of HONO, a precursor of the hydroxyl radical (OH), which represents the primary oxidizing agent in the atmosphere (12).

AOBs oxidize hydroxylamine enzymatically to nitric oxide (NO) by the enzyme hydroxylamine oxidoreductase to extract electrons for ammonia oxidation by the ammonia monooxygenase and respiration (13). Similarly, the hydroxylamine oxidase of the anaerobic ammonium oxidizer *Kuenenia stuttgartienses* detoxifies hydroxylamine to NO and thus generates substrate and electrons for respiration (14). Furthermore, AOB encode a cytochrome P460 (also known as CytL) that detoxifies hydroxylamine by oxidizing it to N_2_O with an unknown biological oxidant (11). So far to the best of our knowledge, the enzymatic conversion of hydroxylamine to N_2_ at physiological conditions has not been reported.

Heterotrophic nitrifiers are able to oxidize ammonia (but cannot grow on it as the only source of energy), and hydroxylamine was identified as an immediate metabolite (15-18). Some of those heterotrophic nitrifiers are also able to denitrify at oxic conditions (19-21), a process called heterotrophic nitrification-aerobic denitrification (HNAD). HNAD has been known for many years, and efforts were made to disclose its metabolic pathway and biochemical mechanism. The general understanding of HNAD is that ammonia is sequentially oxidized to hydroxylamine, NO, nitrite and nitrate by not well understood ammonia-oxidizing enzymes, HAOs and other unknown enzymes. Then, nitrite and nitrate are reduced to N_2_ by the well-known NarGHI/NapAB, NirK/NirS, NorBC and NosZ enzymes (22) that are frequently encoded in the genomes of HNAD bacteria (23, 24). But the understanding of HNAD is still very fragmentary, and both hydroxylamine and nitrite were postulated as links between nitrification and denitrification (6, 21). Furthermore, very little is known about the enzymatic machinery used by heterotrophic nitrifiers for ammonia oxidation (18, 25). In addition, purification and partial characterization of the HAOs from several heterotrophic nitrifiers including *Thiosphera pantotropha* (26), *Pseudomonas* sp. PB16 (27), *Arthrobacter globiformis* (28), and *Acinetobacter* sp. Y16 (29) showed that HAOs from heterotrophic nitrifiers differ significantly from those found in autotrophic nitrifiers, but the functional consequences of these differences have not yet been fully explored.

In this study, we isolated an *Alcaligenes* species strain HO-1, from a wastewater treatment bioreactor. Strain HO-1 was efficiently removing ammonia from wastewater, transiently accumulated hydroxylamine in culture broth, and produced N_2_. Here, we report the genetic cloning and biochemical characterization of a novel oxidase that catalyzed a previously unknown N-N bond-forming reaction from hydroxylamine to N_2_ at physiological conditions.

## Materials and Methods

### Bacterial strains, plasmid, media, cultivation, protein purification and sequence alignment

All bacterial strains, plasmids and primers used in this study are listed in Supplementary Tables S1 and S2. Information about media, cultivation, plasmid construction, protein purification and sequence alignment is provided in Supplemental Information (SI).

### Isolation, characterization, genome sequencing and phylogenetic analysis of strain HO-1

Strain HO-1 was isolated from a SHARON bioreactor (30) using <u>h</u>eterotrophic <u>n</u>itrification <u>m</u>edium (HNM) supplemented with 2.0 mM hydroxylamine, which served as a selective pressure. HNM contained 5 mM (NH_4_)_2_SO_4_ and 17.5 mM succinate, and the concentrations of (NH_4_)_2_SO_4_ and succinate were optimized for fosmid library screening with *E. coli* and transcriptome sequencing with HO-1. The 16S rRNA gene of strain HO-1 was PCR-amplified and extracted also from its sequenced genome. Genome sequencing was performed by Single Molecule, Real-Time (SMRT) technology at the Beijing Novogene Bioinformatics Technology Co., Ltd. Additional information is available as SI.

### Transcriptome sequencing

For transcriptome sequencing (RNA-seq), strain HO-1 was initially cultured in HNM medium containing 14 mM (NH_4_)_2_SO_4_ and 83 mM succinate at 30 °C with shaking as described in SI. After depletion of ammonium at 58 h, 14 mM (NH_4_)_2_SO_4_ were added and cultivation was continued. Samples for RNA extraction were taken at the following time intervals after this (NH_4_)_2_SO_4_ addition, 0 h (C), 3.5 h (T1), 10 h (T2), and 22 h (T3) and then assayed. More detailed information is provided in SI.

### Construction and screening of genomic fosmid library

A genomic fosmid library of strain HO-1 was constructed using the CopyControl™ HTP Fosmid Library Production Kit (Epicentre) according to the instructions of the manufacturer. Fosmid clones were picked in duplicate into two 384-well plates and preserved with 20% glycerol at −80 °C. Clones from one of the duplicate 384-well plates were used as inoculum for four 96-well plates, which contained in each well 160 μl LB with chloramphenicol (12.5 μg/mL). The inoculated plates were cultivated overnight on a high-speed shaker (INFORS HT Microtron, Infors) at 37 °C. 18 μl of the cultured clones were transferred to new 96-well plates, which contained 1.2 mL of HNM with 5 mM (NH_4_)_2_SO_4_, 29.2 mM succinate and 12.5 μg/mL chloramphenicol. Positive clones that were capable of producing hydroxylamine and nitrite from ammonium were screened and sequenced using specific pCC2FOS sequencing primers (Epicentre) at SinoGenoMax (Beijing, China). The genomic locations of inserted DNA fragments were determined by blasting against the genome of strain HO-1.

### Nitrogen-balance experiments

For nitrogen-balance experiments, strain HO-1 or *E. coli* BW25113/pBAD-DnfABC was pre-cultured in HNM with 5 mM (^15^NH_4_)_2_SO_4_ (Spectra Corp., USA) as the sole nitrogen source and transferred three times in this medium for complete depletion of any ^14^N in the cells. Then, 200 μL culture was inoculated into 250 mL sealed bottles containing 20 mL HNM. The headspace of bottles was filled with 1:1 He/O_2_ using a gas displacement system. The bottles were incubated at 30 °C and 160 rpm on a rotary shaker for 80 h (HO-1) or 120 h (*E. coli* BW25113/pBAD-DnfABC). Then, bacterial growth (OD_600_), total nitrogen content (TN), NH_4_ ^+^-N (initial and final amount), gaseous nitrogen products (^15^N_2_, ^15^NO and ^15^N_2_O), soluble nitrogen products (NO_2_^-^-N, NO_3_^-^-N, NH_2_OH-N, organic-N) and biomass-N were measured. Results are presented as mean ± SD from five biological replicates.

### Physiological characterization of recombinant *E. coli* clones

*E. coli* BW25113 carrying pBAD-DnfT1RT2ABCD, pBAD-DnfABC or pBAD-DnfAB was characterized for production of N_2_ and N_2_O, as well as for hydroxylamine accumulation. *E. coli* EPI300™-T1^R^ or *E. coli* BW25113 containing vector pBAD/HisA were used as negative controls under the same conditions. 50 μL of the respective cell suspension with OD_600_ of 0.5 was inoculated into 20 mL sealed tubes containing 5 mL ^15^N-labelled HNM with 0.1% L-arabinose as inducer. Before inoculation, the top air of the sealed tubes was completely replaced by 1 : 1 He/O_2_ as mentioned above. After incubation at 30 °C and 160 rpm for 5 days, gaseous nitrogen compounds (^15^N_2_, ^15^NO, and ^15^N_2_O) and NH_2_OH were detected. Results are presented as mean ± SD from three independent experiments.

### Analytical methods

Bacterial growth was monitored by measuring OD_600_ (spectrophotometer model UV-7200, UNICO, Shanghai, China). Concentrations of ammonium (NH_4_^+^), nitrite (NO_2_^-^) and nitrate (NO_3_^-^) were determined on an Aquakem 600 Analyzer (Thermo Scientific) using the standard methods recommended by the manufacturer. Hydroxylamine was determined by using 8-quinolinol to form the stable 5,8-quinolinequinone-5-(8-hydroxy-5-quinolylimide) (31). Total nitrogen (TN) was determined by using alkaline potassium persulfate digestion with ultraviolet spectrophotometry (32). To determine the biomass nitrogen, bacterial cells for each of the samples were harvested by centrifugation (8 000 *g*, 10 min, 4 °C) and resuspended in ultrapure water with the same volume of the original sample. The nitrogen content of this suspension was determined with the same method as for TN determination, and was calculated as the biomass nitrogen. Quantitative detections of ^15^N_2_ and ^15^NO were performed by GC/MS (model 7890A/5975C, Agilent) equipped with a CP-Molsieve 5A Plot (25 m×0.32 mm×30 μm, Agilent, USA). ^15^N_2_ and NO (which might be formed in the source of mass spectrometers from unlabeled N_2_ and O_2_ and has a similar mass as ^15^N_2_) were eluted at retention times of t=6.897 and t=15.357 min, respectively, and could thus be distinguished without problem (Figure S2). Quantitative detection of ^15^N_2_O was performed as previously described by Liu *et al*. (33).

### ITC measurements

Substrate-enzyme binding experiments were carried out on an ITC200 Isothermal Titration Calorimeter (ITC). All experiments were carried out at 25 °C. The buffer used in this measurement was the same as in the enzymatic assays. 1×0.4 μL followed by 19×2 μL of hydroxylamine were injected into a 200 μL enzyme solution. The data were analyzed using the ORIGIN software (MicroCal Inc.).

### Hydroxylamine oxidase activity assays

*In vitro* enzymatic assays of hydroxylamine oxidase activities were carried out in 300 μL reaction mixtures in buffer (20 mM Tris-HCl, pH 8.5) supplemented with 10 mM ^15^NH_2_OH, 2 mM NADH, 2 mM NADPH and 20 μM FAD. In this experiment, 331 μM DnfA, 3.2 μM DnfB and 311 μM DnfC were added separately or in combination, with mixtures lacking any of these proteins as controls. The reaction was started with the addition of ^15^NH_2_OH (Cambridge Isotope Laboratories Inc.), and the mixture was directly injected into 10 mL gastight tubes, the air of which had been completely replaced by 1:1 He/O_2_. The reactions were incubated at 30 °C without agitation in the dark for 100 min. The hydroxylamine consumption in the mixtures and ^15^N_2_ released in the headspace were assayed. To determinate alternative electron mediators for *in vitro* reconstitution of the hydroxylamine oxidase activity of DnfA, the assay mixture contained 10 μM DnfA, 1 mM ^15^NH_2_OH, 1 mM NADH and various electron mediators including 100 μM flavin adenine dinucleotide (FAD), 100 μM flavin mononucleotide (FMN), 100 μM phenazine methosulfate (PMS), 100 μM phenazine ethosulfate (PES) and 20 μM/10 μM ferredoxin/ferredoxin reductase (Fd/FdR). The reaction mixtures of 300 μL in 10 mL gastight tubes were incubated at 30 °C without agitation in the dark for 60 min. Kinetic assays were conducted in 0.6 μM DnfA, 480 μM FAD, 10 mM NADH and 25-400 μM NH_2_OH of 100 μL reaction mixture in 1.5 mL vials at 30 °C. Enzymatic assays were performed in triplicate.

### Involvement of O_2_ and H_2_ ^18^O measurements

Oxygen (O_2_) requirement experiments were carried out using ^18^O_2_ and detection of the formation of H_2_^18^O. ^18^O_2_/He (with different ratios) was purchased from Newradar Special Gas Co., LTD. (Wuhan, China). The components of 300 μL reaction mixture and reaction conditions were: 331 μM DnfA, 120 μM FAD, 10 mM NADH, 10 mM ^15^NH_2_OH in 20 mM Tris-HCl (pH 8.5), in 10 mL anaerobic tube at different time. The reaction mixtures were diluted 2 times and δ ^18^O was detected by isotopic water analyzer (L2130-I, PICARRO Inc, USA.). The concentration of H_2_^18^O was calculated according to the equation: δ (‰) = (Rsample/Rstandard-1) ×1000. The standard was VSMOW2 (International Atomic Energy Agency; IAEA). Milli-Q water (Millipore, USA) was used as blank controls.

## Results

### *Alcaligenes* sp. strain HO-1 oxidizes ammonia to N_2_ and accumulates hydroxylamine

The bacterial strain HO-1 was isolated from a SHARON bioreactor of a combined system termed “UASB + SHARON + ANAMMOX”, which was used for the treatment of ammonia-rich wastewater (30). Strain HO-1 (electron microscopic pictures of its cell morphology are shown in Figure 1A and 1B) was isolated by using HNM supplemented with 2 mM hydroxylamine as a selective pressure. Purity of strain HO-1 was confirmed by microscopic observation, 16S rRNA gene and genome sequencing. Phylogenomic (Figure 1C) and phylogenetic (supplemental Figure S1) analyses indicated that strain HO-1 and several other *Alcaligenes* isolates formed a coherent cluster, and that this cluster is phylogenetically related but clearly separated from the type strain of *Alcaligenes faecalis*. Average nucleic acid identities (ANI, supplemental Table S3) with all published available genomes of *Alcaligenes* suggest that strain HO-1 belongs to a new, not yet formally described species together with a few *Alcaligenes* strains currently classified as *Alcaligenes faecalis*.

**Figure 1.**
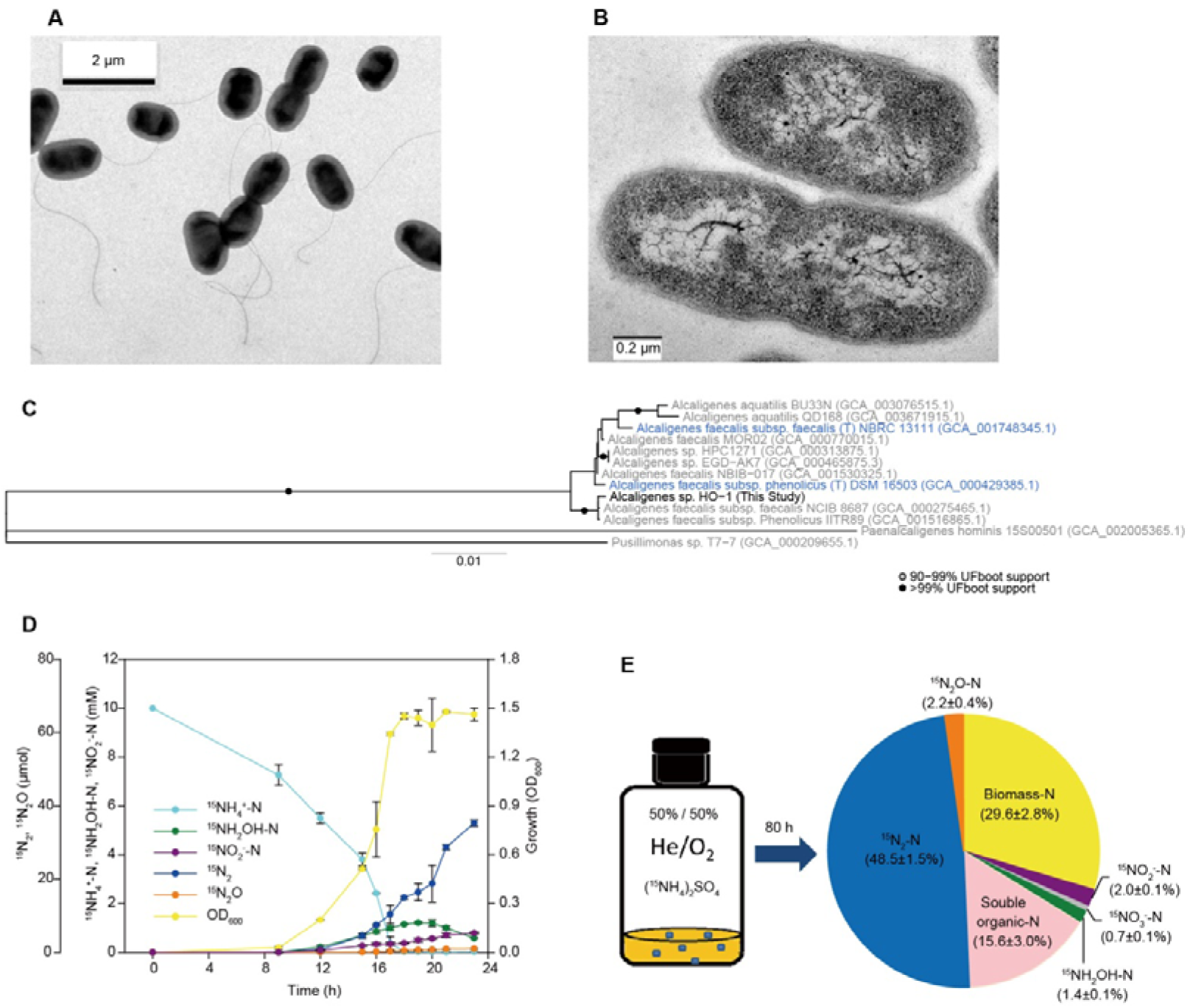
Strain HO-1 converts ammonia mainly into N_2_. (A) and (B) Scanning and transmission electronic microscopic pictures of strain HO-1, respectively. (C) Phylogenomic reconstruction of cultivated *Alcaligenes* with published genomes. *Alcaligenes* sp. HO-1 is depicted in bold and type strains are indicated with a (T) and in blue. Genome accession numbers are given in parentheses. (D) Cell growth (OD_600_), consumption of ^15^NH_4_^+^, transient accumulation of hydroxylamine, and production of ^15^N_2_, ^15^N_2_O, and NO_2_^-^. Production of ^15^NO was not detected. Succinate and (^15^NH_4_)_2_SO_4_ were added at initial concentrations of 17.5 and 5 mM, respectively. (E) The N-balance experiment showed that the consumed ammonia is mainly converted into ^15^N_2_. The data shown in panel (D) and (E) are averages of 3 and 5 replicates, respectively.

Strain HO-1 removed ammonia efficiently under oxic conditions (Table S4) with a tentative accumulation of hydroxylamine in the culture broth (Figure 1D). N-balance experiments of strain HO-1 cultured with 5 mM (^15^NH_4_)_2_SO_4_ showed that all nitrogen atoms were traceable. 48.5% and 2.2% of the consumed ^15^N-labelled ammonium was recovered as ^15^N_2_ and ^15^N_2_O, respectively (Figure 1E), and there was no measurable ^15^NO production (see Figure S2). Small amounts of nitrite (2.0%) and nitrate (0.7%) were detected. Interestingly, ^15^N_2_ production from ammonia by strain HO-1 was not inhibited by 0.1 mM of the copper chelator sodium diethyldithiocarbamate (supplementary Figure S3), which was reported to strongly inhibit at this concentration nitrite reduction to N_2_ in other heterotrophic nitrifiers (24). We also observed that neither nitrate nor nitrite could be used as the sole nitrogen source to support growth of strain HO-1 under oxic conditions (supplementary Figure S4), although nitrite but not nitrate supported anaerobic growth of strain HO-1 and was used as electron acceptor in the presence of succinate as electron donor (supplementary Figure S5). Based on those results, we hypothesized that strain HO-1 might possess an oxic pathway for N_2_ production different from the previously reported aerobic denitrification pathway in HNAD organisms.

### Genome sequence, annotation for N-metabolism genes, and transcriptome analysis of strain HO-1 upon ammonia stimulus

A closed genome of strain HO-1 was determined. Sequence quality control did not find contaminating DNA reads from other organisms, thus further confirming the purity of strain HO-1. The genome of strain HO-1 consists of a single chromosome of 3.77 Mb (Figure 2A). Annotation of N-metabolism related genes (Figure 2B) and their predicted involvements in aerobic ammonia oxidation and anaerobic nitrite reduction are depicted in Supplementary Figure S6. The transcription of these genes was significantly regulated upon ammonia stimulus (Figure 2C). The HO-1 genome encodes *nirK* (encoding a nitrite reductase), *norBC* (encoding nitric oxide reductase) and *nosZ* (encoding nitrous-oxide reductase), representing a complete denitrification pathway from nitrite. This nitrite denitrification pathway is likely responsible for the reduction of nitrite to N_2_O in the presence of O_2_ and to N_2_ in the absence of oxygen with succinate as electron donor, as observed in Figure 2D. This observation is consistent with the perception that nitrous-oxide reductase NosZ is more sensitive to oxygen than other enzymes involved in denitrification and thus might not reduce N_2_O to N_2_ in strain HO-1 under oxic conditions (34). We did not detect any genes coding for an ammonia monooxygenase or hydroxylamine oxidoreductases in the genome of strain HO-1. A pyruvic oxime dioxygenase (POD) is annotated in the genome of strain HO-1, which was previously detected in other *A. faecalis* strains (35). This enzyme catalyzes the oxygenation of pyruvic oxime [2-(hydroxyimino)propanoic acid] to yield pyruvate and nitrite (35-37). In this context it is important to note that hydroxylamine reacts non-enzymatically with pyruvate to pyruvic oxime (38). Consequently, once hydroxylamine is formed from ammonia in this heterotrophic nitrifier, nitrite can be formed in its cytoplasm by a combination of a non-enzymatic reaction and the POD activity (Figure 3C). Consistently, small amounts of nitrite were observed in our ^15^N-ammonium incubation experiments. Besides N_2_O production from abiotic conversion of hydroxylamine (4), partial reduction of nitrite represent a possible origin of N_2_O in the N-balance experiments. We did not find any *nrf* and *nap* genes in the genome of strain HO-1, which are indicative for the absence of dissimilatory or assimilatory nitrate reduction to ammonia. This is consistent with the observation that strain HO-1 did not use nitrate as nitrogen source for growth. The small amounts of nitrate detected in the ammonium addition experiments might be formed by nitrite-oxidizing side-activity of the strain HO-1 catalases KatA and KatE (39, 40) (Figure 3C). Alternatively, nitrate formation might have been catalyzed by the NO detoxifying enzyme, flavohemoglobin-NO-dioxidase (Hmp), which converts NO together with O_2_ to NO_3_^-^ (41) as strain HO-1 has two genes with significant homologies to *hmp*.

**Figure 2.**
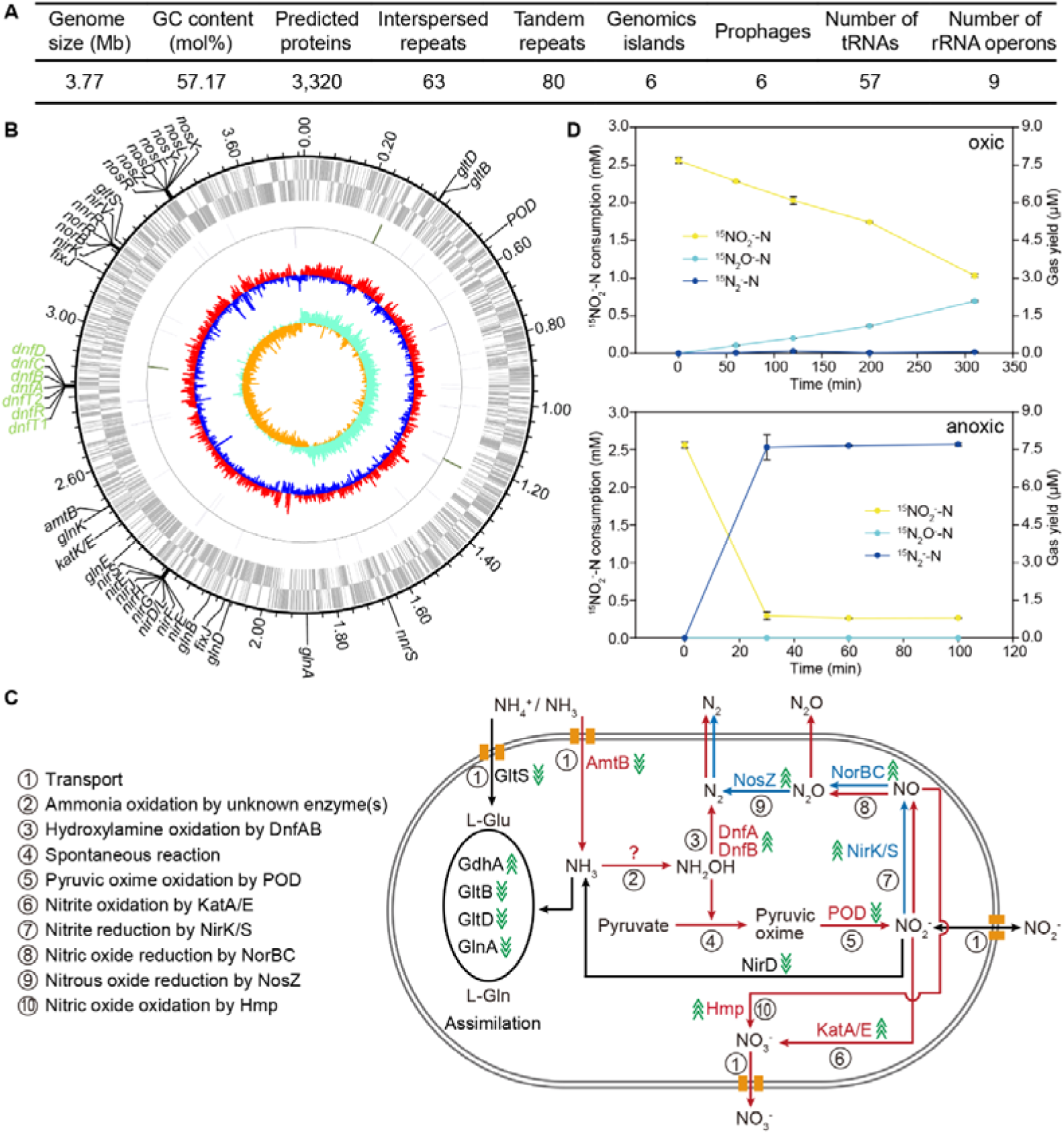
Panel A summarizes general features of the genome of strain HO-1. Panel B shows a circular genome map with predicted N-metabolism-related genes. Circles from the outermost to the center: (1) scale marks of the genome, (2) protein-coding genes on the forward strand, (3) protein-coding genes on the reverse strand, (4) tRNA (purple) and rRNA (dark green) genes on the forward strand, (5) tRNA (purple) and rRNA (dark green) genes on the reverse strand, (6) GC content, (7) GC skew. Panel C shows a schematic representation of the proposed routes for N-metabolism in strain HO-1 based on gene annotation and transcriptomic data (green arrows; for details see Table S5). The routes for aerobic ammonia oxidation are showed in red, and anaerobic denitrification is showed in blue. Panel D shows experimental date of nitrite conversion by strain HO-1 in HNM medium to N_2_O at oxic or to N_2_ at anoxic conditions. The data shown in panel (D) are averages of 3 replicates.

**Figure 3.**
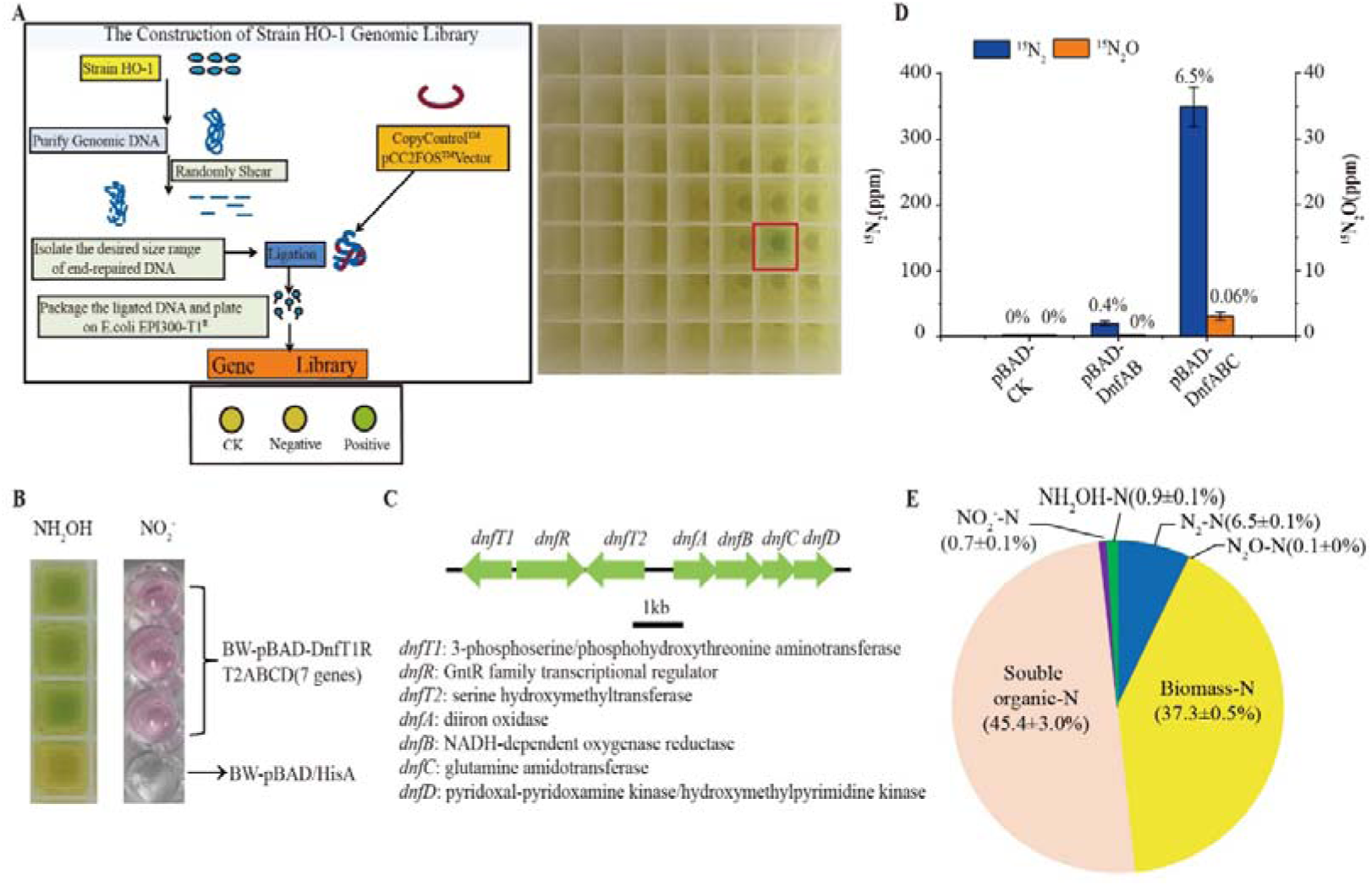
*dnfABC* genes from strain HO-1 enable recombinant *E. coli* ammonia conversion into N_2_. (A) Flowchart summarizing the construction of the fosmid library and the screen for hydroxylamine accumulation. On the right side, one of the six positive clones (see text) that accumulated hydroxylamine is spotted (framed in red). Ck, a control assay performed using *E. coli* EPI300™-T1^R^ that did not harbor any exogenous gene. (B) Demonstration for hydroxylamine and nitrite production in *E. coli* harboring gene cluster *dnfT1RT2ABCD*. (C) Annotation of the *dnf* genes are shown on the right side, and their homolog proteins in UniProtKB/Swiss-Prot database are listed in Table S6. (D) Production of N_2_ and N_2_O by *E. coli* clones harboring *dnfABC* or *dnfAB* genes. (E) ^15^N-balance results from *E. coli* cells carrying *dnfABC*. The numbers indicate the ratio (%) of ^15^N_2_ or ^15^N_2_O production to the consumption of ammonia. All experiments were conducted under 50% O_2_ + 50% He in HNM medium with 17.5 mM succinate and 5 mM (^15^NH_4_)_2_SO_4_ in 10 mL gastight tubes and 250 mL sealed bottles. The data shown in panel (D) and (E) are averages of 3 and 5 replicates, respectively.

In order to identify genes that are involved in ammonia oxidation and N_2_ formation and to check whether the above discussed N-metabolism genes were regulated in strain HO-1, we performed stimuli-response transcriptome experiments. Strain HO-1 was cultured with 83 mM succinate and 14 mM of ammonium sulfate until ammonia was completely consumed and then 14 mM of ammonium sulfate was added to the culture. Cells were harvested at 0 h, 3.5 h, 10 h and 22 h after the addition of ammonium sulfate, and were subjected to transcriptomic analysis. Supplemental Table S5 show the transcriptional response of N-metabolism-related genes to this ammonia stimulus. The transcription levels of genes involved in denitrification (*nirK, norBC, nosZ*) and glutamate metabolism (*gltD, gltB, gltS, glnK*) were significantly (fold changes >2) regulated. As discussed above, the POD and KatA activities might catalyze the formation of nitrite and nitrate, respectively, and we found their transcripts upregulated at 22 h upon ammonia stimulus. We also found that the *amtB* (encoding a putative ammonia transporter), *gltD, gltB, gltS*, and *glnK* were significantly downregulated, which were reasonable responses to the high concentration of ammonia. Interestingly, we observed that a gene without a functional annotation (FE795_13025, named *dnfA*) and its neighboring genes (FE795_13030, FE795_13035, FE795_13040; named *dnfB, dnfC*, and *dnfD*, respectively) were all significantly upregulated at 3.5 h and 10 h in response to ammonium stimulation, and their transcriptional levels decreased as ammonium was depleted at 22 h. As shown in Figure 2C (inside panel), the transcriptional levels of *dnfA* and *dnfB* were very high after 3.5 h.

### The gene cluster *dnfABC* from strain HO-1 enables *E. coli* to accumulate hydroxylamine and to produce N_2_ from ammonia

In order to identify the pathway and genes involved in the aerobic ammonia oxidation and N_2_ production in strain HO-1, we constructed a fosmid library of the strain HO-1 genome in *E. coli* strain EPI300™-T1^R^ (Figure 3A). We screened 1920 clones of the fosmid library, and six clones that were positive for hydroxylamine accumulation were obtained. Sequencing the 6 clones revealed that they shared a cluster of 7 genes (named *dnfT1RT2ABCD, dnf* stands for di***n***itrogen ***f***ormation) (Figure 3C), which were also detected on the chromosome of strain HO-1 (their position is shown in Figure 2B). As described above, two (*dnfA,dnfB*) of the 7 genes were significantly upregulated and strongly transcribed upon ammonia stimulation. Excitingly, these 7 genes conferred *E. coli* cells the ability to produce hydroxylamine and nitrite (Figure 3B), N_2_ and N_2_O from ammonium. We subcloned the *dnfABC or dnfAB* genes into *E. coli* strain BW25113 cells. *E. coli* cells carrying *dnfABC or dnfAB* already generated N_2_ and N_2_O from added ammonium, although *E. coli*/pBAD*-*DnfAB showed much decreased N_2_ production when compared to *E. coli*/pBAD-DnfABC (Figure 3D). In N-balance experiments of *E. coli* harboring *dnfABC* cultured with HNM media, 6.5% of the consumed ^15^N-labelling were recovered as ^15^N_2_ (Figure 3E).

The interpretation of data from physiological experiments with a host carrying heterologously expressed genes obviously also has to take into account the relevant pathways naturally encoded by the host cells. Both the *E. coli* strain BW25113 (for subcloning) and strain BL21(DE3) (for heterologously expressing proteins, see below) encodes in their genomes genes for nitrate transport (Nrt), dissimilatory nitrate reduction (NarGHI, NapAB, NirBD, NrfAH), nitrate reduction/nitrite oxidation (NxrAB) and assimilation (supplementary Figure S6). This *E. coli* strain encodes no enzymes that would catalyze N_2_ and NH_2_OH production via nitrification and/or denitrification pathways. Thus, the observed production of N_2_ and NH_2_OH from ^15^NH_4_^+^ in the *E. coli* strain carrying *dnfABC* from strain HO-1 cannot be explained by the metabolism of *E. coli*. We therefore concluded that *dnfABC* is critical to catalyze the conversion of ammonia to N_2_ in the *E. coli* cells.

### DnfA binds and converts hydroxylamine into N_2_

Based on our bioinformatic analyses, DnfA, DnfB and DnfC were annotated as diiron oxygenase, NADH-dependent oxygenase reductase, and glutamine amidotransferase, respectively. *In silico* analyses of DnfA, DnfB and DnfC did not reveal transmembrane domains or secretion signals, and thus most likely these proteins have a cytoplasmic localization. DnfB has binding sites for NAD(P)H, FMN and 2Fe-2S. Since the genetic cluster *dnfABC* conferred *E. coli* cells the ability to produce hydroxylamine and N_2_ from ammonia, firstly, we individually cloned and expressed *dnfA, dnfB*, and *dnfC* in *E. coli*. The DnfA, DnfB and DnfC were purified to homogeneity (Figure 4A) and their interactions with ammonia and hydroxylamine were assayed with an isothermal titration calorimeter (ITC). Results showed that none of them interacted with ammonia, but DnfA bound to hydroxylamine with a dissociation constant (Kd) of 0.16 ± 0.02 μM (Figure 4B). Next, we tested whether DnfA, DnfB, or DnfC was able to catalyze the conversion of ammonia and hydroxylamine. An *in vitro* reconstitution enzymatic activity assay using the chemical electron mediator FAD was performed following the previously reported method for the DnfA homolog AurF (42). We found that neither DnfA, nor DnfB, nor DnfC catalyzed the oxidation of ammonia *in vitro*, but that DnfA, as well as the combinations of DnfA/DnfB and DnfA/DnfB/DnfC, catalyzed the conversion of hydroxylamine to N_2_ in the presence of molecular O_2_, FAD and NADH (Figure 4C). We calculated that the consumption of approximate 2 moles of hydroxylamine produced one mole N_2_, suggesting hydroxylamine was almost stoichiometrically converted into N_2_. Considering that hydroxylamine was completely converted to ^15^N_2_ when DnfA was present but not so when DnfA was absent, we concluded that DnfA was essential. It was reported that chemical decomposition of hydroxylamine in soils produced N_2_ (43). We observed trace but detectable amount of N_2_ production in the reaction system without DnfA. A possible explanation might be that O_2_ was activated by FAD/NADH and consequently triggered spontaneous reactions with yet unknown chemistry for N_2_ generation. But the amount of N_2_ production was much lower than the one observed from the DnfA-catalyzed conversion of hydroxylamine (Figure 4C), indicating a major role of DnfA catalysis.

**Figure 4.**
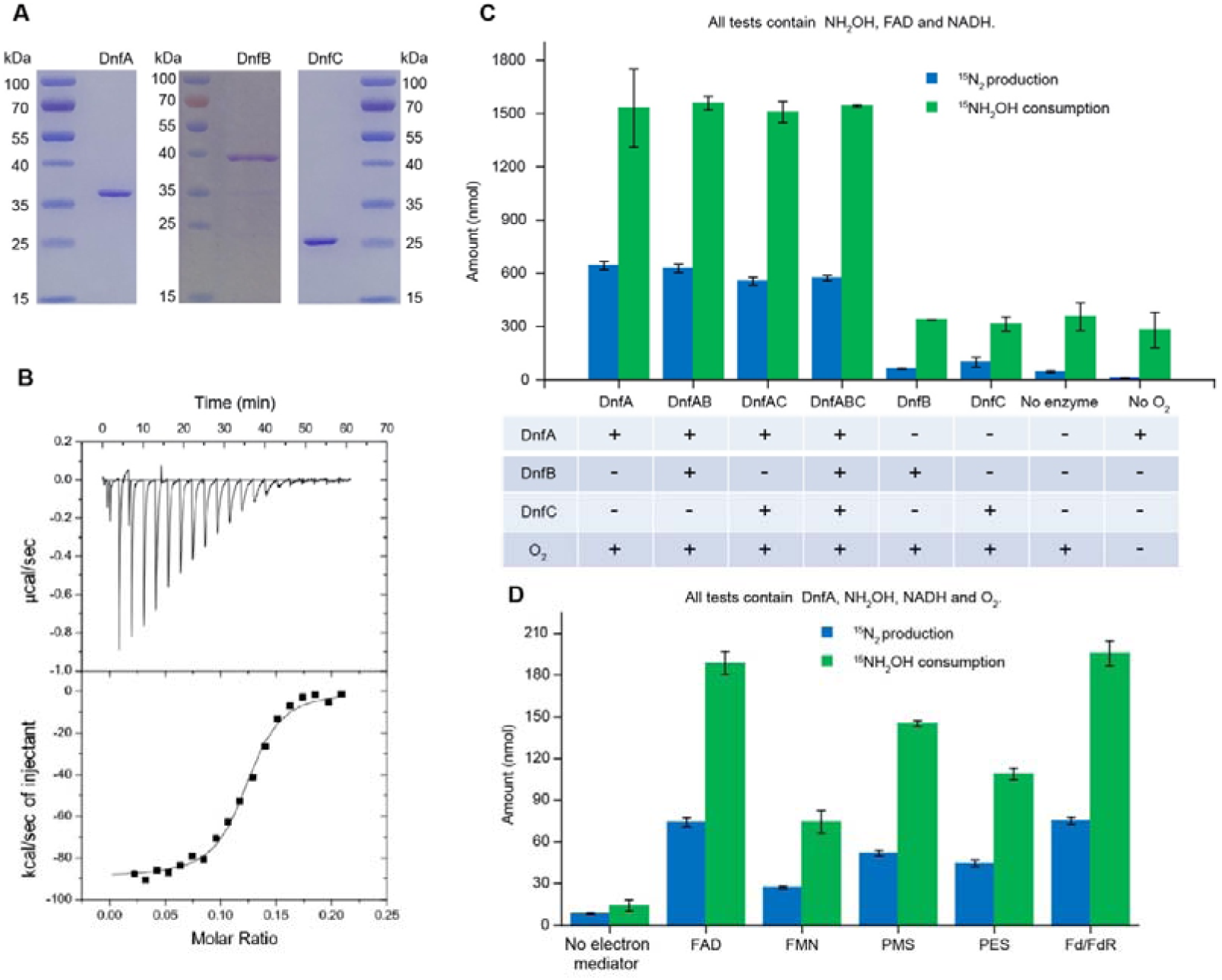
DnfA binds to and catalyzes the conversion of hydroxylamine to N_2_. (A) SDS-PAGE of purified DnfA, DnfB and DnfC. (B) ITC assay of the DnfA-hydroxylamine interaction. 0.2 mM hydroxylamine was titrated into 0.1 mM DnfA. The experiments were repeated three times, and a representative result is shown. (C) Enzymatic assay of DnfA activity (331 μM DnfA, 3.2 μM DnfB and 311 μM DnfC) in presence of 10 mM ^15^NH_2_OH, 20 μM FAD, 2 mM NADH or 2 mM NADPH and 50% O_2_. (D) Determination of suitable electron mediators for reconstitution of hydroxylamine oxidase activity of DnfA *in vitro*. The assays were performed with 10 μM DnfA, 1 mM hydroxylamine, 1 mM NADH and various electron mediators, including FAD (100 μM), FMN (100 μM), phenazine methosulfate (PMS, 100 μM), phenazine ethosulfate (PES, 100 μM) and ferredoxin/ferredoxin reductase (Fd/FdR, 20 μM and 10 μM). Data shown in panels C and D are averages of 3 replicates.

DnfB contained both ferredoxin (Fd) and ferredoxin reductase (FdR) domains and its gene is located at the immediate vicinity of DnfA. Thus, it seems likely that DnfB is an Fd/Fd reductase and mediates electron transfer to DnfA. This is consistent with our observation that Fd/FdR/NADH supported DnfA activity during *in vitro* assays. In addition, we also found that additional electron donors such as phenazine methosulfate and phenazine ethosulfate also supported DnfA activity in the presence of NADH (Figure 4D).

### DnfA is a hydroxylamine oxidase

DnfA contains a diiron-motif and was annotated to be a diiron oxygenase. We determined that iron atoms were associated with DnfA molecules at a ratio of 1.60 ± 0.01 : 1, suggesting that one molecule of DnfA bound two iron atoms. DnfA homologues were identified in the genomes of *Burkholderia, Delftia, Herbaspirillum, Microvirgula, Pantoea*, and *Pseudomonas* species, in addition to *Alcaligenes* species. Those DnfA homologues formed a coherent clade and they are related to the AurF N-oxygenase (42, 44) and CmlI homologues (Figure 5A), and is distantly related to previously known HAOs (34, 45, 46) (sequence identity < 20% over full length). We demonstrated that DnfA activity depends on molecular oxygen by performing experiments with ^18^O_2_ (Figure 4C). By using 10 mM ^15^N-labelled hydroxylamine and 10% of ^18^O_2_ (90% of the gas phase is Helium), we determined the molar ratio (net reaction catalyzed by DnfA) of hydroxylamine consumption to N_2_ production to H_2_^18^O production was 2.6 : 1 : 1.9 (Figure 5B). We also calculated this ratio with hydroxylamine consumption rate, ^15^N_2_ and H ^18^O production rates under the same condition. The consumption rate of hydroxylamine was determined to be 8.71 ± 1.29 μM/min per mg protein, and the production rates of ^15^N_2_ and H ^18^O were 3.84 ± 0.64 μM/min per mg protein and 7.53 ± 0.64 μM/min per mg protein, respectively (Figure 5C). The molar ratio of hydroxylamine consumption rate to N_2_ production rate to H_2_^18^O production rate was calculated to be 2.3:1:1.9, a ratio close to the values in equation (1). Similar assays were carried out under 1% ^18^O_2_ and the result also confirmed the equivalent consumption/production rates of NH_2_OH, N_2_ and H_2_O. Further, we determined the kinetic constants of DnfA-catalyzed reactions at atmospheric condition (21% O_2_). The Michaelis-Menten constant (Km) for hydroxylamine and the catalytic efficiency (kcat) were determined to be 48.1 ± 6.0 μM and 18.6 ± 1.2 min^-1^, respectively (Figure 5D). The substrate binding affinity and catalytic efficiency of DnfA are comparable with those reported for the DnfA homolog AurF (Km 5.24 ± 0.64 μM and kcat 6.21 ± 0.52 min^-1^), supporting the possibility of hydroxylamine as the substrate under physiological conditions. Based on the above results, a model was proposed for the DnfA-catalyzed oxidation of hydroxylamine to N_2_ (Figure 5E).

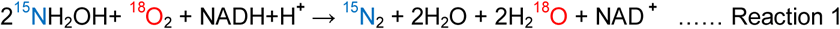

**Figure 5.**
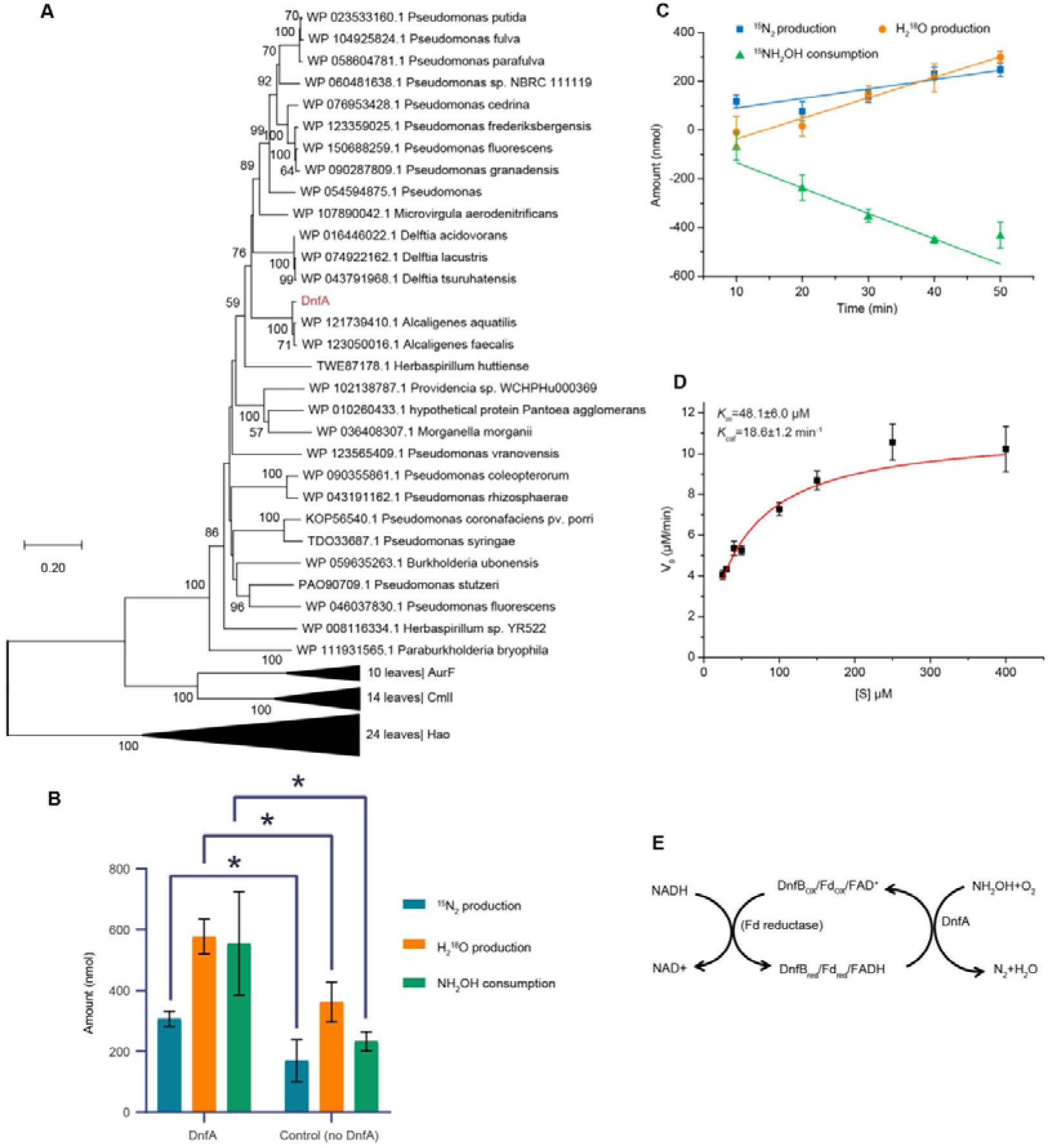
Isotope-labelling experiment with (^15^NH_4_)_2_SO_4_ and ^18^O_2_ showing the involvement of O_2_ during hydroxylamine oxidation and determination of kinetic constants of DnfA. (A) Neighbour-joining phylogenetic tree of DnfA, AurF, CmlI and HAO, and their homologues. Bootstrap percentages based on 1,000 replicates are shown at branch points. AurF, p-aminobenzoate N-oxidase from *Streptomyces thioluteus*. CmlI, N-Oxidase from *Streptomyces venezuelae*. DnfA, AurF and CmlI are indicated by asterisks. The tree showing DnfA, AurF and CmlI fell within the same linage, and HAOs formed a distinct linage from them. 24 HAOs leaves were collapsed. (B) The consumption or production of NH_2_OH, ^15^N_2,_ and H_2_^18^O with (left) or without enzyme (right) after 20 min incubation. Experiments were performed in 0.3 ml reaction solution with initial 10 mM ^15^NH_2_OH, 120 μM FAD, 10 mM NADH, 331 μM DnfA, and 10% of ^18^O_2_ + 90% of He in the head gas space. The asterisk (*) represents statistically significant (P<0.05). (C) Consumption rate of NH_2_OH and production rates of ^15^N_2_ and H_2_^18^O. Experiments were performed with the same protocol as in (B). The data shown here are the differences of the amounts of substrates and products from triplicate experiments with and without DnfA, and data are fitted into a linear equation to access the reaction rates. (D) Determination of kinetic constants of DnfA under atmospheric conditions. The assays were performed with 0.6 μM DnfA, 480 μM FAD, 10 mM NADH and 25-400 μM NH_2_OH (100 μL reaction mixture in 1.5 mL vials at 30 °C). For kinetic constant determination at another temperature including lower NH_2_OH concentrations see Fig. S7 (E) The proposed catalytic model of DnfA. The data shown here are averages of 3 replicates.

## Discussion

While a reasonable albeit not complete understanding of the enzymes and pathways involved in aerobic autotrophic ammonia oxidation is available, our knowledge on the physiology and biochemistry of heterotrophic nitrification is surprisingly rudimentary (23, 47). In this study, we discovered in the newly isolated heterotrophic nitrifier *Alcaligenes* strain HO-1 a previously unknown process that leads to the oxidation of ammonia to N_2_ gas. Via heterologous expression in *E. coli* and work with purified Dnf proteins/enzymes, we demonstrated that the gene cluster *dnfT1T2RABCD* is critical for this process. While we did not yet manage to reveal which enzyme(s) is catalyzing the conversion of ammonia to hydroxylamine in the *Alcaligenes* strain HO-1, we demonstrated that the key enzyme DnfA catalyzes a new reaction - the oxidation of hydroxylamine and NADH with oxygen to N_2_ and water [reaction 1]. It should be noted that oxygen can be non-enzymatically reduced to water by NADH (Figure 5B), but our data demonstrated that the oxidation of hydroxylamine clearly relied on O_2_. Additional research is urgently required to reveal the biochemical mechanism of this novel way of N-N bond formation.

An obvious key question is which physiological role the oxidation of ammonia to N_2_ via hydroxylamine plays in *Alcaligenes* HO-1. In the reaction catalyzed by DnfA [reaction 1], two electrons are needed for the formation of one molecule N_2_ and four molecules of water from two molecules of hydroxylamine and one molecule of oxygen. In addition, ammonia oxidation to hydroxylamine (if catalyzed by an unknown AMO) costs additional 2 electrons. Thus, it is tempting to speculate that *Alcaligenes* uses this reaction to flare off electrons during rapid growth and that this process serves to maintain redox balance in this organism. The DnfA catalyzed process will also help to reduce the hydroxylamine concentration in the cytoplasm and thus minimize chemical conversion of precious pyruvate to pyruvic oxime (which subsequently needs to be salvaged by the activity of the POD enzyme). However, clearly additional physiological experiments will be needed to investigate this in more detail.

The discovery of hydroxylamine oxidation with an oxidase to N_2_ in an *Alcaligenes* strain is particularly noteworthy as several *Alcaligenes* strains were previously reported as heterotrophic nitrifiers producing N_2_ gas (6, 16, 20, 21, 48, 49). In this context, it is interesting to note that homologues of *dnfA* are found in other *Alcaligenes* species as well as in other heterotrophic nitrifiers such as *Pseudomonas, Microvirgula, Pokkaliibacter*, and *Burkholderia*, suggesting that this hydroxylamine oxidase may widely occur in diverse microorganisms. If this reaction indeed is responsible under aerobic conditions for (part of the) N_2_-formation from ammonia in many heterotrophic nitrifiers, our discovery will fundamentally change our perception on heterotrophic nitrification and might offer applications in engineered systems for N-removal from contaminated waters.

## Acknowledgments

We thank Dr. Sebastian Luecker at Department of Microbiology, Radboud University, the Netherlands, and Prof. Holger Daims at University of Vienna for helpful discussions during the preparation of this manuscript. This research was supported by grants from National Key R&D Program of China (2019YFA0905500; 2016YFD0501409) and National Natural Science Foundation of China (NSFC grant Nos. 31861133002; 91951101; 31870103), Chinese Academy of Science (KFZD-SW-219-3; KFZD-SW-309), and CAS Key Technology Talent Program (to GMA).

## Data and materials availability

The genome sequence of *Alcaligenes* sp. strain HO-1 has been deposited at NCBI GenBank under the BioProject PRJNA543690 with the accession number CP049362.

## Author Contributions

M-RW, YL, Y-XZ, T-TH isolated the strain HO-1, conducted the N-balance, constructed the fosmid library and finished the genetic cloning and screening; L-LM, M-RW, LM, H-ZZ, X-YG, Y-LQ, T-TH, TW, G-MA finished protein purification, enzymatic assays, biochemical tests and isotope measurements; S-JL, Z-PL, D-FL and X-HS designed and supervised the experiments; M-RW, T-TH, H-ZZ, CWH, S-JL, Z-PL, and D-FL analyzed the data; and MW, Z-PL, D-FL and S-JL wrote the manuscript.

