## Supplemental Information for "A novel oxidase from *Alcaligenes* sp. HO-1 oxidizes hydroxylamine to N_2_"

ISME Journal

Running Title: A novel oxidase catalyzes hydroxylamine oxidation to N_2_

**Other Supplementary Materials for this manuscript include the following:**

- Supplementary Figures S1-S7.
- Supplementary tables S1-S6.
- Dataset S1 RNA-seq assays of strain HO-1 stimulated by ammonia.

**Bacterial strains, plasmid, media, and cultivation**

All bacterial strains and plasmids used in this study are listed in Table S1. Luria–Bertani medium (LB) and modified heterotrophic nitriﬁcation medium (HNM) were prepared according to references (1). Modified HNM contained 5 mM (NH_4_)_2_SO_4_ and 17.5 mM succinate unless otherwise stated. *Alcaligenes* sp. strain HO-1 was routinely cultivated in modified HNM media, aerobically at 30 °C. For denitrification tests, 5 mM nitrite or nitrate was added as electron acceptor. Growth of strain HO-1 under anaerobic conditions was tested in an anaerobic jar with an AnaeroPack (Mitsubishi Gas Chemical Company Inc., Tokyo, Japan), and growth was observed for one week. *E. coli* was cultivated at 37 °C in LB media. For preparation of agar plates, 1.5% (w/v) of agar was added to the media. Ampicillin (100 µg/mL), or chloramphenicol (12.5 µg/mL) was supplemented to media as necessary.

**Phylogenetic analysis of strain HO-1**

The 16S rRNA gene of strain HO-1 was PCR-amplified and extracted also from sequenced genome. Reference sequences for related type and non-type strains were downloaded from the Ribosomal Database Project (2). Downloaded sequences were aligned using mafft-linsi (3) and gap-filtered (setting -gt 0.95) using trimAl (4). Phylogenetic reconstruction was calculated in IQTREE (5) with model HKY+I+G4 and 1,000 ultrafast bootstraps. To construct the phylogenomic tree, genomes were downloaded from GenBank for all available cultured strains of *Alcaligenes*, *Paenalcaligenes* and *Pusillimonas* strains with published genomes. An alignment of 34 concatenated marker genes was generated for downloaded genomes using CheckM (6) and gap-filtered (setting -nogaps) using trimAl (4). Phylogenomic reconstruction was calculated in IQTREE (5) with model LG+F+G4 and 1,000 ultrafast bootstraps.

**Genome sequencing**

Genomic DNA of strain HO-1 was extracted using a TIANamp Bacterial DNA Kit (Tiangen Biotech, Beijing, China). The extracted DNA was checked for quality with agarose gel electrophoresis and quantified by Qubit. Genome sequencing was performed by Single Molecule, Real-Time (SMRT) technology at the Beijing Novogene Bioinformatics Technology Co., Ltd.. Briefly, DNA sample qualified by electrophoresis was used to construct a 10 Kb SMRT Bell library followed by sequencing using the PacBio RSII platform. Raw data were trimmed by the SMRT 2.3.0 (7, 8), and filtered reads were assembled with the SMRT portal generating one contig without gaps. Open reading frames were predicted using GeneMarkS (9). Average amino acid identity (AAI) and average nucleotide identity (ANI) were calculated with a nonredundant set of putative genes predicted by Prodigal (10) as well as annotated genes. gANI was calculated using Microbial Species Identifier (MiSI) (11).

**RNA extraction and transcriptome sequencing**

For transcriptome sequencing (RNA-seq), strain HO-1 was initially cultured in HNM medium containing 14 mM (NH_4_)_2_SO_4_ and 83 mM succinate at 30 °C with shaking as described above. After incubation for 58 h (when ammonium was depleted), the culture was amended again with (NH_4_)_2_SO_4_ (final concentration, 14 mM) and cultivation was continued. Samples for RNA extraction were taken at the following time intervals after (NH_4_)_2_SO_4_ addition, 0 h (C), 3.5 h (T1), 10 h (T2), and 22 h (T3). Cells from samples were immediately collected by centrifuging (8,000 rpm, 2 min, 4 °C) and were frozen with liquid nitrogen and stored at -80 °C until used for RNA extraction. Total RNA was extracted using the E.Z.N.A.TM Bacterial RNA Kit (Omega, Norcross, GA, USA). Genomic DNA was removed with RNase-free DNase I (TaKaRa, Shiga, Japan). PCR targeting RecA gene fragment using primer pair recA-F/R was performed to check for any DNA contamination. Purified RNA was quantified by Nanodrop 2000. RNA quality was detected by agarose gel electrophoresis and an Agilent 2100 Bioanalyzer system (Agilent Technologies, Inc., Santa Clara, CA, USA). Three biological replicates were carried out.

Two μg of total RNA of each sample were used for Illumina RNA-seq with a strand-specific RNA sequencing method (12, 13). Ribosomal RNA was depleted by subtractive hybridization using Ribo-Zero Magnetic kit (Epicentre, Madison, Wisconsin, USA). Paired-end RNA-seq sequencing libraries were constructed by TruSeqTM RNA sample preparation Kit (Illumina, San Diego, CA, USA). Sequencing was performed on an Illumina HiSeq×10 (2×150 bp read length) at Shanghai Majorbio Pharm Technology Co. Ltd. (Shanghai, China). The sequence data from the Illumina platform were analyzed on the free online platform of Majorbio I-Sanger Cloud Platform (www.i-sanger.com). Gene transcription levels were quantified by RSEM (14) and reported as TPM values (15, 16). Normalization and differential expression analysis was performed by DESeq2 (17). Multiple testing of false discovery rate (p-adjust) was calculated using Benjamini/Hochberg method (18).

**Plasmid construction**

Primers used for PCR amplification were listed in Table S1. Genomic DNA of *Alcaligenes* sp. HO-1 and *E. coli* strains were prepared as mentioned above. Plasmid DNA was prepared with the TIANprep Mini Plasmid Kit (Tiangen Biotech). PCR amplification was performed using the TransStart® FastPfu DNA Polymerase (Transgen, Beijing, China). Purification of DNA fragments from the PCR reaction was performed using the Gel Extraction Kit (Omega, Norcross, GA, USA) or Cycle Pure Kit (Omega). DNA fragments were ligated into linearized vectors using the ClonExpress II One Step Cloning Kit (Vazyme, Nanjing, China). Plasmid pBAD-DnfT1RT2ABCD was constructed from the pBAD/HisA vector (Amp^r^; Invitrogen, Carlsbad, USA). Linear pBAD/HisA with the deletion of *araBAD* promoter was amplified using primers pBAD-VF/pBAD-VR. Two fragments covering *dnfT1RT2ABCD* were amplified with primer pairs seq1-F/seq1-R and seq2-F/seq2-R from the genomic DNA of strain HO-1. These fragments were each introduced an overlapping area at both ends by PCR and were joined via Gibson assembly (19) to produce plasmid pBAD-DnfT1RT2ABCD using the ClonExpress II One Step Cloning Kit (Vazyme). To construct pBAD-DnfABC and pBAD-DnfAB, the pBAD/HisA vector was linearized with primers pBAD-F/pBAD-R. Segments of *dnfABC* and *dnfAB* were respectively amplified from strain HO-1 using the primer pairs as listed in Table S2. To construct pET-21a-DnfA and pET-21a-DnfB, pET-21a was linearized with primer pair pET21a-F/pET21a-R, the *dnfA* and *dnfB* was respectively amplified from strain HO-1 using the primer pairs as listed in Table S1. After circularization, *dnfA* or *dnfB* was inserted into pET-21a (+) vector with a 6xHis-tag located at C-terminus. To construct pBAD-DnfC for expressing DnfC with a 6xHis-tag located at C-terminus, linear pBAD/HisA was obtained by primer pair pBAD-hisF/pBAD-R, dnfC was amplified with primer pair dnfC-F/dnfC-R, then they were circularized to generate plasmid pBAD-DnfC.

**Purification of DnfA, DnfB, DnfC and ferredoxin reductase**

To prepare proteins DnfA, DnfB and DnfC, *E. coli* BL21(DE3) cells carrying the plasmid pET-21a-DnfA or pET-21a-DnfB and *E. coli* BW25113 carrying the plasmid pBAD-DnfC were grown in LB supplied with 100 µg/mL ampicillin at 37 °C and 200 rpm. When the OD_600_ reached 0.3~0.6, *E. coli*/pET-21a-DnfA and *E. coli*/pET-21a-DnfB were induced with 0.5 mM IPTG at 16 °C on a rotary shaker (160 rpm) for 20 h, and *E. coli*/pBAD-dnfC was induced with 0.1% L-arabinose at 16 °C overnight. The cells were harvested by centrifuging at 5,000 *g* for 30 min at 4 °C, resuspended in buffer A (100 mM Tris-HCl, 100 mM NaCl, 10 mM imidazole, pH 8.0), and subsequently lysed by ultrasonication with 99 cycles (work for 4 s and pause for 4 s by each cycle) at 200 W. After centrifugation at 14,000 *g* for 30 min at 4 °C and filtered through 0.45 μM filter, the supernatant was applied to a Ni-NTA resin (Qiagen) column which was previously equilibrated with buffer A. Then the Ni-NTA matrix was washed with buffer A by adding imidazole at a 10–50 mM concentration to remove impurities. 6xHis-tagged protein was eluted with buffer B (100 mM Tris-HCl, 100 mM NaCl, 250 mM imidazole, pH 8.0). The purified 6xHis-tagged proteins were desalted using centrifugal filter devices (Merck Millipore) and exchanged into buffer C (20 mM Tris-HCl, pH 8.5) with a PD-10 desalting column (GE Healthcare, USA). Ferredoxin (selFdx1499, Fd) and ferredoxin reductase (selFdR0978, FdR) were expressed according to the previously reported method and purified following the method described above (23). Protein concentrations were determined by Bradford method using a Quick Start Protein Assay Kit (Bio-Rad, USA).

**Multiple sequence alignment of DnfA homologues**

The amino acid sequences of DnfA, AurF (CAE02601.1) and CmlI (WP_015032130.1) were as query to perform blastp against the nr database with expect threshold ≤ 0.001 in NCBI, respectively. Filter results at a ≥ 90% sequence identity level and ≥ 60% query coverage, and choose only one representative sequence from each species. All homolog sequences of HAO were downloaded from K10535 orthology in KEGG (20) with one representative selected in each species. Representative sequences from *Streptomyces thioluteus* HKI-227 (AJ575648.1, AurF), *Alcaligenes aquatilis* (WP_121739410.1), *Alcaligenes faecalis* (WP_123050016.1), *Delftia acidovorans* (WP_016446022.1), *Microvirgula aerodenitrifica* (WP_107890042.1), *Pseudomonas* sp. (WP_054594875.1), *Herbaspirillum* sp. YR522 (WP_008116334.1), and *Streptomyces venezuelae* ATCC 10712 (WP_015032130.1, CmlI) were downloaded from NCBI (https://www.ncbi.nlm.nih.gov/). The multiple sequence alignment was performed using Clustal Omega program (21). To construct phylogenetic tree of DnfA and related proteins, MEGA7 software (22) for Windows was used to align multiple sequences with Muscle algorithms (23). Protein distances were calculated by applying a neighbor-joining method. 1,000 bootstrap resampling were analyzed to test phylogeny.

**Gas production analysis of denitrification**

To identify the gas production of denitrification, HO-1 cells were harvested, resuspended to 3 mL ammonia and succinate depleted HNM (OD_600_ of 10) that contained 2 mM ^15^N-labelled nitrite, and then transferred into 250 mL sealed bottles. The headspace of bottles was filled with 1 : 1 He/O2 (50% O2) using a gas displacement system. The production of ^15^N_2_ and ^15^N_2_O and the consumption of nitrite were monitored.

**Inhibition of copper nitrite reductase in HO-1 strain**

To identify whether diethyldithiocarbamate (DDT) works in HO-1 cells, the nitrite consumption by HO-1 cells was assayed at different concentration of DDT. Strain HO-1 cells of early-exponential-phase were harvested by centrifugation (12, 000 g, 5 min, 30 °C) after shake flask culturing in HNM aerobically at 30 °C. Cells were washed three times and resuspended to OD_600_ of 10 with ammonia and succinate depleted HNM that contains 2 mM nitrite and 0, 0.05, 0.1, 0.4, or 1 mM DDT, culturing into Erlenmeyer flasks on a rotary shaker at 30 °C and 160 rpm. The concentration of nitrite was monitored after incubating for 30, 80, 130, 200, 300 and 360 min.

To assay the effect of DDT on gas production, strain HO-1 cells were treated with and without DDT. As described above, cells were harvested, resuspended to OD_600_ of 10, and transferred into 120 mL sealed bottles containing 10 mL ^15^N-labelled HNM (5 mM (^15^NH_4_)_2_SO_4_ (Spectra Corp., USA)) with or without 0.1 mM DDT filled 50% O2 and 50% He. The bottles were incubated on a rotary shaker at 30 °C and 160 rpm. ^15^N_2_ and ^15^N_2_O were measured after incubating for 30, 70, 110, 150 and 190 min.

**Table S1 Bacterial strains and plasmids used in this study**

| Strain/plasmid | Description | Source/reference | |
| --- | --- | --- | --- |
| Strains |  | |  |
| *Alcaligenes* sp. HO-1 | Converting ammonia to N_2_ | | This study |
| *E. coli* strains |  | |  |
| EPI300™-T1^R^ | Fosmid cloning host | | Epicentre |
| BL21(DE3) | Protein expression host | | Transgen |
| BW25113 | Protein expression host | | Invitrogen |
| Plasmids |  | |  |
| pCC2FOS | Fosmid cloning vector | | Epicentre |
| pBAD/HisA | Gene expression vector | | Invitrogen |
| pBAD-DnfT1RT2ABCD | pBAD/HisA carrying *dnfT1RT2ABCD* gene cluster (FE795_13010 to FE795_13040) with araBAD promoter deleted from the vector | | This study |
| pBAD-DnfABC | pBAD/HisA carrying *dnfABC* (FE795_13025 to FE795_13035) | | This study |
| pBAD-DnfAB | pBAD/HisA carrying *dnfAB* (FE795_13025 to FE795_13030) | | This study |
| pBAD-DnfC | pBAD/HisA carrying *dnfC* (FE795_13035) | | This study |
| pET-21a (+) | Gene expression vector | | Novagen |
| pET-21a-DnfA | pET-21a (+) carrying *dnfA* (FE795_13025) | | This study. |
| pET-21a-DnfB | pET-21a (+) carrying *dnfB* (FE795_13030) | | This study. |
| pET-28a-Fd | pET-28a vector carrying [2Fe-2S] ferredoxin (Fd) | | (24) |
| pET-28a-FdR | pET-28a vector carrying ferredoxin-NADP^+^ reductase (FdR) | | (24) |

**Table S2** **Primers used in this study**

| Name | Sequence (5’→3’) | Description |
| --- | --- | --- |
| recA-F | CGCCGCTCTGTCGCAAAT | For PCR targeting RecA gene fragment |
| recA -R | ACGCCCAAAGCAATGTCC |  |
| pBAD-VF | AGCTTGGCTGTTTTGGCGGATGAGA | For PCR of a linar fragment from the circular pBAD for cloning of *dnfT1RT2ABCD* |
| pBAD-VR | GGTCCCGCTTTGTTACAGAATGCTT |  |
| seq1-F | TTCTGTAACAAAGCGGGACCTCAAATCGCAGCCACCTGCC | For PCR of the former fragment of *dnfT1RT2ABCD* (*dnfT1RT2A*) |
| seq1-R | ATTACCTGAGCCGATCCGTTCTG |  |
| seq2-F | AACGGATCGGCTCAGGTAATTCT | For PCR of the later fragment of *dnfT1RT2ABCD* (*dnfABCD*) |
| seq2-R | TCCGCCAAAACAGCCAAGCTCAAACAGCATGGCCATCCAGG |  |
| pBAD-F | AAGCTTGGCTGTTTTGGCGG | For PCR of a linar fragment from the circular pBAD for cloning of *dnf* genes. |
| pBAD-R | GGTTAATTCCTCCTGTTAGCCCAAAAA |  |
| dnfABC-F | GGGCTAACAGGAGGAATTAACCATGACTATCAAAAGCTACGAAACCGATGA | For PCR of *dnfABC* |
| dnfABC-R | CCGCCAAAACAGCCAAGCTTTCATGCAGCACAATCGGCTTT |  |
| dnfAB-F | GGGCTAACAGGAGGAATTAACCATGACTATCAAAAGCTACGAAACCGATGA | For PCR of *dnfAB* |
| dnfAB-R | CCGCCAAAACAGCCAAGCTTTCAGCTCGTGGTGCCGAT |  |
| pET21a-F | AAGCTTGCGGCCGCACTCG | For PCR of a linar fragment from the circular pET21a |
| pET21a-R | GGATCCGCGACCCATTTGCTGTC |  |
| pet-dnfA-F | GCAAATGGGTCGCGGATCCATGACTATCAAAAGCTACGAAACCG | For PCR of *dnfA* |
| pet-dnfA-R | TCGAGTGCGGCCGCAAGCTTTTGCAGCGCCTCCTGTTG |  |
| pet-dnfB-F | GCAAATGGGTCGCGGATCCATGACAGCCATGATTCAGGCG | For PCR of *dnfB* |
| pet-dnfB-R | CGAGTGCGGCCGCAAGCTTGCTCGTGGTGCCGATGTCTA |  |
| pBAD-hisF | CCACCACCACCACCACTGAAAGCTTGGCTGTTTTGGCGG | For PCR of a linar fragment from the circular pBAD/HisA to construct pBAD-DnfC |
| pBAD-hisR | GGTTAATTCCTCCTGTTAGCCCAAAAA |  |
| dnfC-F | GCTAACAGGAGGAATTAACCATGAAGAAAGTCATCGCACTGCG | For PCR of *dnfC* |
| dnfC-R | TCAGTGGTGGTGGTGGTGGTGTGCAGCACAATCGGCTTTGT |  |

**Table S3 Average nucleotide identity (ANI) of strain HO-1 with related *Alcaligenes faecalis* (A.f.) strains**

| **Organisms** | **Same species to A.f.?** | **accession** | **sources** | **gANI** |  |  | **gANI_cov** |
| --- | --- | --- | --- | --- | --- | --- | --- |
| *A. faecalis* subsp. phenolicus IITR89 | Y | GCA_001516865.1 | 26941148 | 99.4 |  |  | 0.9 |
| *A. faecalis* subsp. faecalis NCIB 8687 | Y | GCA_000275465.1 | 22933773 | 99.3 |  |  | 0.905 |
| *A. faecalis* subsp. phenolicus (T) DSM 16503 | N | GCA_000429385.1 | 25197443 | 86.9 |  |  | 0.8 |
| *A. faecalis* subsp. faecalis (T) NBRC 13111 | N | GCA_001748345.1 | 28059164 | 86.9 |  |  | 0.835 |
| *Alcaligenes* sp. HPC1271 | N | GCA_000313875.1 | 23469352 | 86.7 |  |  | 0.75 |
| *Alcaligenes* sp. EGD-AK7 | N | GCA_000465875.3 | 24407646 | 86.7 |  |  | 0.805 |
| *A. faecalis* MOR02 | N | GCA_000770015.1 | 25540337 | 86.6 |  |  | 0.795 |
| *A. faecalis* NBIB-017 | N | GCA_001530325.1 | 27056227 | 86.6 |  |  | 0.81 |
| *A. aquatilis* QD168 | N | GCA_003671915.1 | 30714040 | 86.3 |  |  | 0.785 |
| *A. aquatilis* BU33N | N | GCA_003076515.1 | 31550268 | 86.1 |  |  | 0.83 |
| *Pusillimonas* sp. T7-7 | N | GCA_000209655.1 | 21622753 | 72.7 |  |  | 0.355 |
| *Paenalcaligenes hominis* 15S00501 | N | GCA_002005365.1 | 28450518 | 70.45 |  |  | 0.24 |

Note: Type strains are indicated with (T) in the organism name. Typically, organisms with >96.5% gANI and >0.6 coverage can be considered the same species. Genome accessions and literature sources are provided. Source is given as pubmed ID numers.

**Table S4 Growth of and ammonium removal by strain HO-1 with different initial ammonium concentrations under aerobic conditions**

| Initial NH_4_^+^ (mM) | Culture time (h) | OD_600_ | Residue NH_4_^+^  (mM) | Removal  Rates (%) |
| --- | --- | --- | --- | --- |
| 10 | 18 | 0.81±0.01 | 0.05±0.00 | 99.50±0.00 |
| 28 | 48 | 2.48±0.03 | 0.06±0.00 | 99.79±0.00 |
| 57 | 144 | 5.18±0.19 | 0.82±0.01 | 98.56±0.02 |
| 86 | 192 | 8.42±0.07 | 15.29±0.74 | 82.22±0.86 |

Succinate was used as sole carbon source with a C/N mass ratio of 10 : 1. For the assays with 10, 28 and 57 mM ammonia, the data shown here were those at the time when ammonia was almost depleted (in a continuous monitor of the ammonia concentration). For the assay with 86 mM ammonia, the data shown here were those after cultivation for 192 hours. All results were from triplicate experiments.

**Table S5** **Transcriptions of genes involved in denitrification and glutamate metabolism upon addition of ammonium sulfate in the presence of succinate**. Significant fold changes are highlighted in yellow.

| Gene ID |  | TPM | | | | Log2FC (p-adjust) | | |
| --- | --- | --- | --- | --- | --- | --- | --- | --- |
|  |  | 0h | 3.5h | 10h | 22h | 3.5h/0h | 10h/0h | 22h/0h |
| FE795_00995 | *gdhA* | 79.05±14.13 | 167.71±6.36 | 134.22±3.42 | 68.52±13.20 | -0.12 (2.12E-01) | 0.07 (5.33E-01) | 0.11 (3.45E-01) |
| FE795_01695 | *gltD* | 196.84±61.67 | 92.61±36.92 | 143.59±25.54 | 22.31±6.67 | -2.27 (3.85E-19) | -1.14 (1.32E-13) | -2.83 (5.37E-79) |
| FE795_01700 | *gltB* | 483.02±100.28 | 138.20±53.11 | 190.27±32.07 | 37.20±10.58 | -2.99 (2.38E-36) | -2.04 (9.04E-56) | -3.39 (1.03E-190) |
| FE795_02445 | *POD* | 326.12±152.61 | 213.65±18.34 | 173.09±10.60 | 1169.03±141.54 | -1.80 (4.58E-19) | -1.57 (6.37E-16) | 2.21  (8.37E-27) |
| FE795_06380 | *gltS* | 20.24±1.47 | 22.52±2.97 | 16.21±1.83 | 6.70±1.03 | -1.08 (5.85E-12) | -1.05 (5.51E-12) | -1.29 (1.92E-12) |
| FE795_07360 | *hmp* | 0.00±0.00 | 0.42±0.37 | 0.48±0.31 | 0.09±0.15 | 0 (1.00E+00) | 0 (1.00E+00) | 0 (1.00E+00) |
| FE795_07530 | *nnrS* | 149.89±28.61 | 266.82±23.35 | 334.69±33.09 | 83.09±19.22 | -0.38 (3.71E-04) | 0.46 (1.61E-06) | -0.54 (1.00E-05) |
| FE795_08655 | *glnA* | 705.67±61.21 | 783.50±94.28 | 669.41±91.80 | 184.69±42.09 | -1.07 (4.18E-18) | -0.79 (5.59E-12) | -1.64 (4.14E-50) |
| FE795_09645 | *glnD* | 13.26±1.63 | 23.29±1.72 | 18.74±1.03 | 2.60±0.66 | -0.40 (1.21E-03) | -0.21 (6.41E-02) | -2.05 (3.93E-51) |
| FE795_09700 | *fixJ* | 63.66±10.52 | 120.62±4.51 | 93.00±6.18 | 32.70±6.07 | -0.31 (4.85E-03) | -0.16 (1.58E-01) | -0.65 (2.43E-07) |
| FE795_09885 | *glnB* | 57.65±12.92 | 96.42±4.79 | 83.08±9.70 | 36.98±12.95 | -0.52 (1.57E-04) | -0.17 (2.41E-01) | -0.36 (3.79E-02) |
| FE795_10470 | *nirE* | 1.84±0.88 | 1.39±0.50 | 2.41±0.16 | 0.66±0.39 | -1.63 (4.75E-04) | -0.28 (5.05E-01) | -1.20 (2.47E-02) |
| FE795_10480 | *nirF* | 0.83±0.29 | 2.13±0.76 | 2.15±0.49 | 0.43±0.19 | 0.19 (7.00E-01) | 0.70 (6.43E-02) | -0.68 (1.77E-01) |
| FE795_10485 | *nirD/L* | 1.04±0.12 | 2.21±0.39 | 2.33±0.25 | 0.80±0.22 | -0.14 (7.37E-01) | 0.45 (2.07E-01) | -0.08 (8.75E-01) |
| FE795_10490 | *nirG* | 0.20±0.18 | 2.05±0.39 | 1.34±0.18 | 0.11±0.12 | 2.12 (2.60E-02) | 2.06 (3.43E-02) | -0.68 (7.30E-01) |
| FE795_10495 | *nirH* | 1.67±1.39 | 3.22±1.15 | 2.87±0.20 | 0.63±0.41 | -0.26 (6.75E-01) | 0.12 (8.37E-01) | -1.11 (1.50E-01) |
| FE795_10500 | *nirJ* | 2.08±0.46 | 4.01±0.09 | 3.63±0.18 | 2.44±0.07 | -0.26 (3.43E-01) | 0.11 (6.77E-01) | 0.58 (3.72E-02) |
| FE795_10505 | *nirE* | 2.30±0.90 | 8.51±1.36 | 6.67±0.15 | 3.48±0.88 | 0.68 (6.34E-03) | 0.84 (2.68E-04) | 0.91 (3.44E-04) |
| FE795_10515 | *nirS* | 1.72±0.36 | 10.83±2.67 | 3.15±0.22 | 1.45±0.14 | 1.45 (1.37E-10) | 0.18 (4.58E-01) | 0.09 (7.46E-01) |
| FE795_10925 | *gdhA* | 12.20±1.99 | 61.59±4.79 | 27.02±0.80 | 2.81±0.57 | 1.12 (1.54E-22) | 0.45 (7.25E-04) | -1.81 (3.10E-29) |
| FE795_10985 | *glnE* | 12.13±3.68 | 32.71±1.92 | 26.55±2.23 | 6.26±1.60 | 0.25 (1.04E-01) | 0.44 (9.58E-04) | -0.63 (3.27E-05) |
| FE795_11315 | *katE* | 2.28±0.49 | 11.43±0.44 | 7.13±±0.45 | 3.25±0.64 | 1.12 (1.11E-12) | 0.94 (4.33E-09) | 0.84 (2.34E-05) |
| FE795_11520 | *glnK* | 366.94±72.30 | 139.46±51.59 | 224.88±39.02 | 58.09±7.61 | -2.62 (9.56E-31) | -1.41 (3.23E-30) | -2.34 (5.06E-96) |
| FE795_11525 | *amtB* | 147.43±25.91 | 28.77±4.33 | 51.35±4.89 | 6.86±2.04 | -3.57 (5.77E-165) | -2.22 (2.70E-106) | -4.12 (7.13E-164) |
| FE795_14375 | *fixJ* | 2.17±0.43 | 5.53±1.07 | 5.90±0.96 | 2.08±0.75 | 0.12 (7.48E-01) | 0.74 (1.34E-02) | 0.22 (5.58E-01) |
| FE795_14730 | *nirK* | 10.37±12.93 | 527.88±181.10 | 149.82±39.85 | 2.69±0.22 | 4.60 (5.61E-06) | 3.30 (4.83E-04) | -1.45 (1.44E-01) |
| FE795_14735 | *norZ* | 4.24±1.49 | 331.01±265.42 | 15.08±4.79 | 3.28±0.64 | 5.09 (1.63E-07) | 1.16 (3.76E-06) | -0.03 (8.75E-01) |
| FE795_14740 | *norR* | 12.18±1.40 | 17.18±2.37 | 13.45±0.56 | 8.61±2.81 | -0.73 (1.74E-07) | -0.56 (2.16E-05) | -0.22 (1.88E-01) |
| FE795_14745 | *nnrR* | 19.83±2.24 | 41.60±3.32 | 45.12±3.73 | 16.90±7.86 | -0.16 (2.48E-01) | 0.48 (1.20E-03) | 0.03 (9.11E-01) |
| FE795_14750 | *nirV* | 1.08±0.69 | 82.47±24.03 | 28.33±7.13 | 1.12±0.09 | 5.09 (2.71E-75) | 4.05 (2.40E-52) | 0.45 (2.03E-01) |
| FE795_15610 | *nosR* | 0.40±0.49 | 41.68±15.57 | 5.79±1.90 | 0.28±0.05 | 5.63 (1.38E-09) | 3.28 (4.76E-04) | -0.01 (9.89E-01) |
| FE795_15615 | *nosZ* | 0.26±0.05 | 307.48±181.69 | 22.43±4.83 | 0.34±0.05 | 8.98 (2.66E-38) | 5.75 (1.26E-72) | 0.70 (1.67E-01) |
| FE795_15620 | *nosD* | 0.36±0.12 | 101.06±59.57 | 8.37±2.61 | 0.71±0.11 | 6.93 (5.23E-23) | 3.87 (6.50E-23) | 1.33 (6.18E-03) |
| FE795_15625 | *nosF* | 0.16±0.13 | 44.28±27.79 | 3.92±1.22 | 0.34±0.09 | 6.94 (1.67E-13) | 3.94 (1.28E-10) | 1.48 (9.67E-02) |
| FE795_15630 | *nosY* | 0.15±0.17 | 22.27±13.02 | 2.37±1.11 | 0.31±0.17 | 6.03 (1.33E-07) | 3.35 (7.75E-06) | 1.36 (2.07E-01) |
| FE795_15635 | *nosL* | 0.07±0.12 | 34.88±19.79 | 2.94±1.46 | 0.42±0.47 | 7.81 (6.83E-09) | 4.74 (8.47E-05) | 2.87 (8.73E-02) |
| FE795_15640 | *nosX* | 1.93±0.73 | 64.34±36.75 | 9.50±0.56 | 4.83±0.66 | 3.86 (1.94E-10) | 1.60 (1.19E-12) | 1.66 (1.21E-10) |
| FE795_16420 | *hmp* | 1.24±0.49 | 8.60±1.59 | 4.26±0.19 | 2.23±0.84 | 1.59 (3.34E-09) | 1.09 (4.47E-05) | 1.14 (1.89E-04) |
| FE795_17180 | *katA* | 47.78±15.71 | 92.78±4.73 | 113.59±7.95 | 323.38±7.61 | -0.23 (1.58E-01) | 0.57 (7.86E-05) | 3.13 (4.60E-65) |
| FE795_13010 | *dnfT1* | 188.69±31.44 | 346.83±25.19 | 165.36±26.54 | 94.74±17.48 | -0.34 (1.49E-04) | -0.89 (5.31E-10) | -0.68 (3.29E-12) |
| FE795_13015 | *dnfT2* | 25.38±1.19 | 40.60±5.13 | 34.30±2.74 | 33.92±9.85 | -0.55 (3.86E-05) | -0.29 (2.37E-02) | 0.70 (3.40E-06) |
| FE795_13020 | *dnfR* | 13.00±2.43 | 36.53±1.19 | 22.90±1.44 | 19.02±3.29 | 0.29 (1.52E-02) | 0.12 (3.43E-01) | 0.87 (6.52E-13) |
| FE795_13025 | *dnfA* | 1011.70±259.06 | 6518.63±411.38 | 4908.24±364.15 | 790.14±175.82 | 1.49 (7.39E-32) | 1.59 (1.23E-40) | -0.04 (7.94E-01) |
| FE795_13030 | *dnfB* | 413.41±88.39 | 4038.26±492.94 | 2907.45±341.37 | 353.28±108.63 | 2.08 (2.76E-56) | 2.12 (1.11E-75) | 0.07 (6.20E-01) |
| FE795_13035 | *dnfC* | 359.78±75.73 | 3509.55±381.29 | 2491.89±243.62 | 280.01±75.34 | 2.06 (2.34E-63) | 2.09 (6.72E-89) | -0.06 (6.42E-01) |
| FE795_13040 | *dnfD* | 26.45±4.15 | 448.40±45.63 | 242.90±25.92 | 13.56±4.10 | 2.85 (5.73E-86) | 2.49 (5.45E-62) | -0.68 (9.29E-05) |
| FE795_10260 | *recA* | 329.47±25.44 | 315.04±32.86 | 194.47±21.88 | 225.09±15.43 | -1.30 (2.82E-27) | -1.48 (9.63E-35) | -0.22 (1.85E-01) |

**Table S6** **Homolog proteins of *dnf* genes found in UniProtKB/Swiss-Prot database**

| Query Protein | Description | Target Organisms | Query Coverage | Identity | Target Accession |
| --- | --- | --- | --- | --- | --- |
| DnfT1 | phosphoserine/phosphohydroxythreonine transaminase | *Acinetobacter baumannii* ACICU | 99% | 46.80% | B2HWW3.1 |
| DnfR | Probable rhizopine catabolism regulatory protein MocR | *Sinorhizobium meliloti* | 95% | 33.61% | P49309.1 |
| DnfT2 | serine hydroxymethyltransferase/methylase | *Bordetella bronchiseptica* RB50 | 95% | 60.05% | Q7WPH6.1 |
| DnfA | Non-heme di-iron N-oxygenase | *Streptomyces venezuelae* ATCC 10712 | 84% | 27.78% | F2RB83.1 |
| DnfB | NADH oxidoreductase HCR reducing the hybrid cluster protein HCP (a high-affnity NO reductase and hydroxylamine reductase) | *Escherichia coli* K-12 | 85% | 24.42% | P75824.3 |
|  | Phthalate dioxygenase reductase | *Burkholderia cepacia* | 89% | 24.05% | P33164.3 |
|  | Toluene-4-sulfonate monooxygenase system reductase subunit TsaB1/TS methylmonooxygenase system, reductase B | *Comamonas testosteroni* | 88% | 27.62% | P94680.1 |
| DnfC | Putative glutamine amidotransferase-like protein YfeJ | *Salmonella enterica* subsp. enterica serovar Typhimurium str. LT2 | 73% | 34.81% | P40194.2 |
|  | Putative glutamine amidotransferase-like protein L716 | *Acanthamoeba polyphaga mimivirus* | 90% | 25.00% | Q5UNX1.1 |
| DnfD | Pyridoxine/pyridoxal/pyridoxamine/B6-vitamer kinase | *Bordetella parapertussis* 12822 | 95% | 49.27% | Q7W6K7.2 |

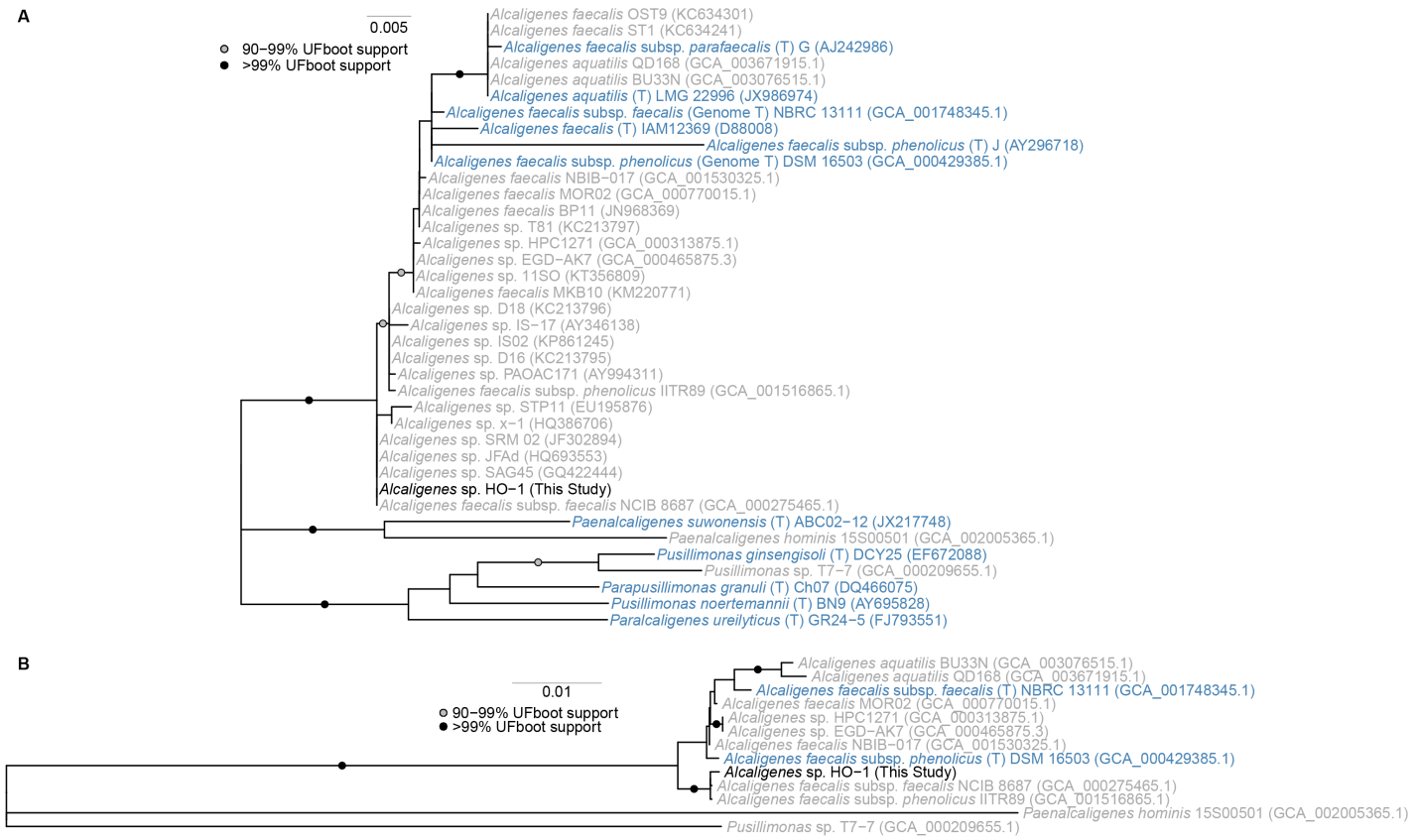

**Fig. S1. Neighbour-joint phylogenetic tree of strain HO-1 and its closely related isolates within the genus *Alcaligenes*, based on 16S rRNA gene sequences.** *Alcaligenes* sp. HO-1 is labeled black. Type strains are in blue text and indicated with (T). Data are from publicly available 16S rRNA gene sequences and and labeld with T for those obtained from publicly available genomes. The 16S rRNA gene sequence accession numbers or genome assembly accession numbers are given in parentheses.

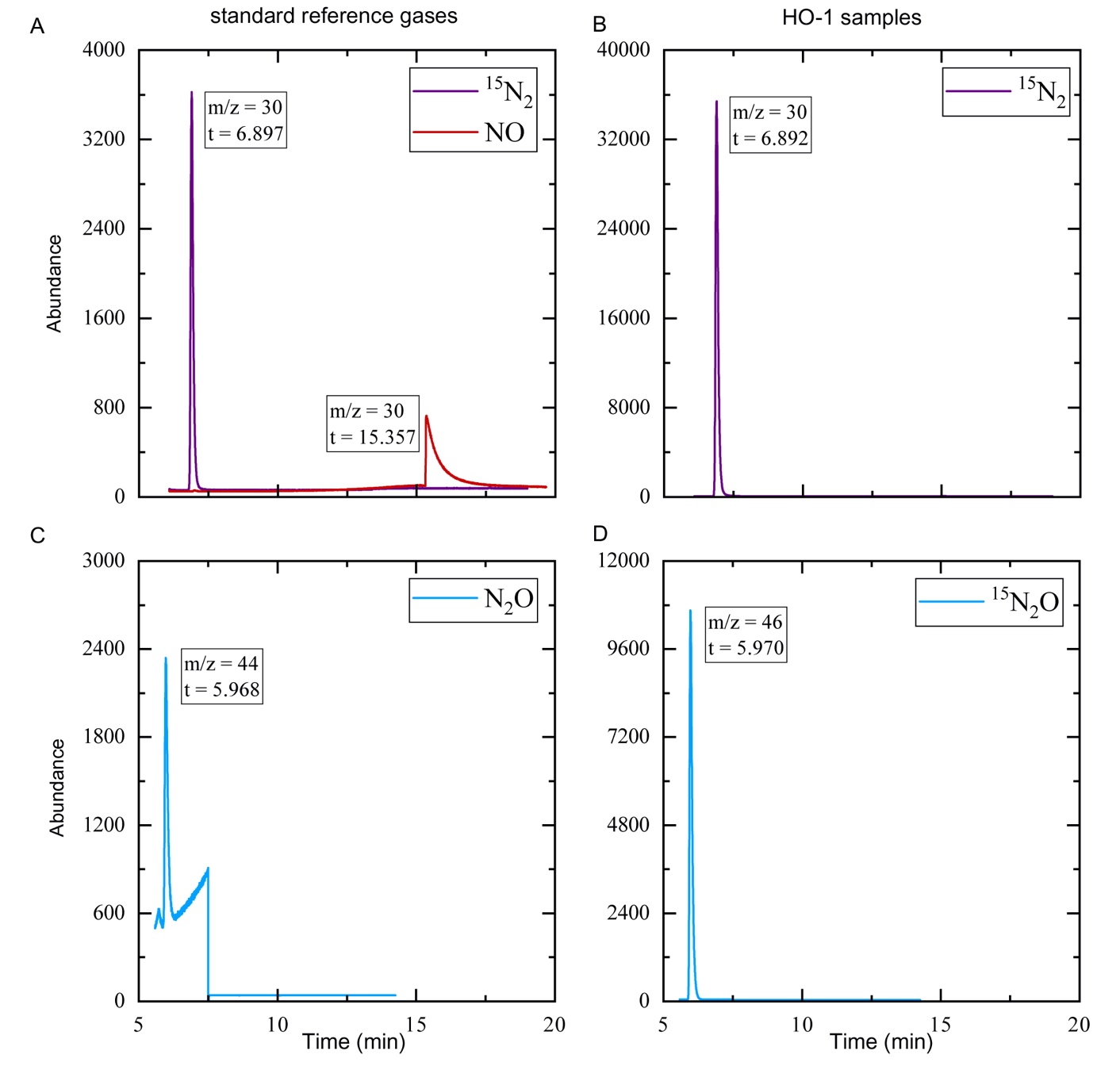

**Fig. S2. Assay of ^15^N_2_, ^15^N_2_O and ^15^NO with GC-MS.** Standard reference ^15^N_2_, NO (A) and N_2_O gases (C) and HO-1 sample gases (B and D) were loaded on a CP-Molsieve 5A Plot column (panels A and B, ^15^N_2_ and NO) and/or a GS-Carbon Plot column (panels C and D, N_2_O). The relative abundances of gases with their m/z values and retention times are shown.

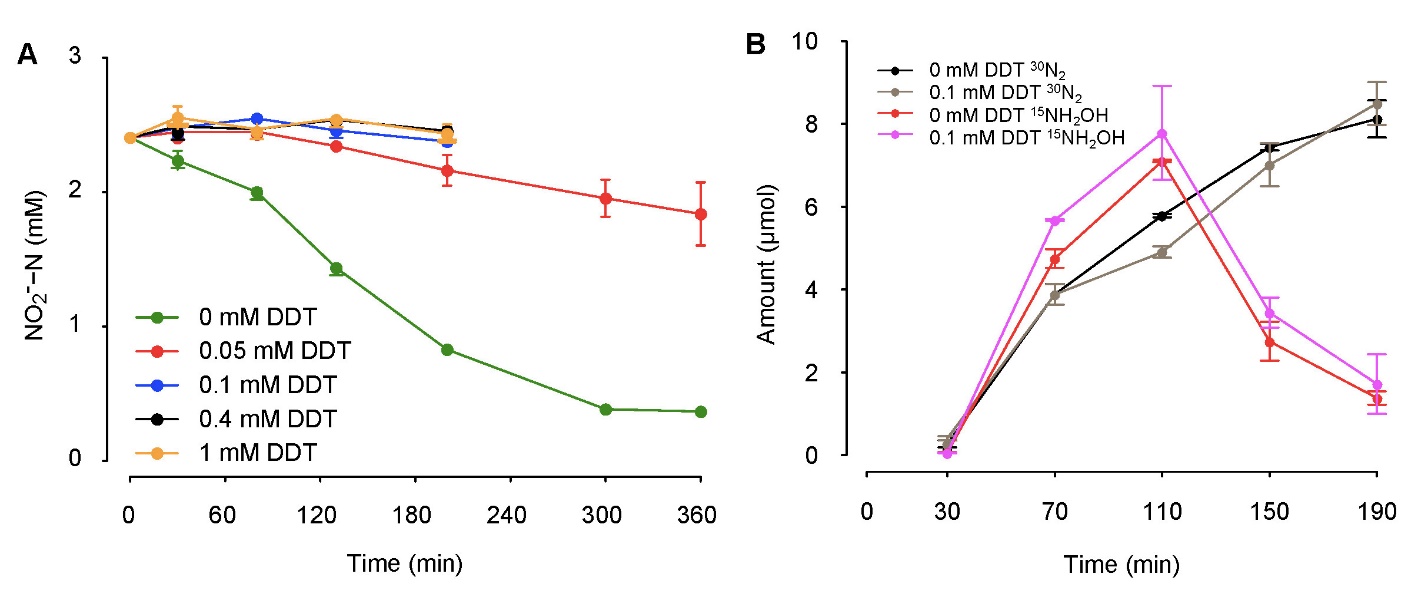

**Fig. S3.** Inhibition of copper nitrite reductase of strain HO-1 by sodium diethyldithiocarbamate (DDT). (A) Strain HO-1 cells were harvested, resuspended to OD_600_ of 10, and incubated in the presence of 2 mM nitrite with/without DDT at atmospheric condition. The residual nitrite was monitored. (B) The production of ^15^N_2_ and ^15^NH_2_OH from (^15^NH_4_)_2_SO_4_ were not obviously affected in the presence of 0.1 mM diethyldithiocarbamate (DDT). HO-1 cells were cultured in HNM with or without DDT

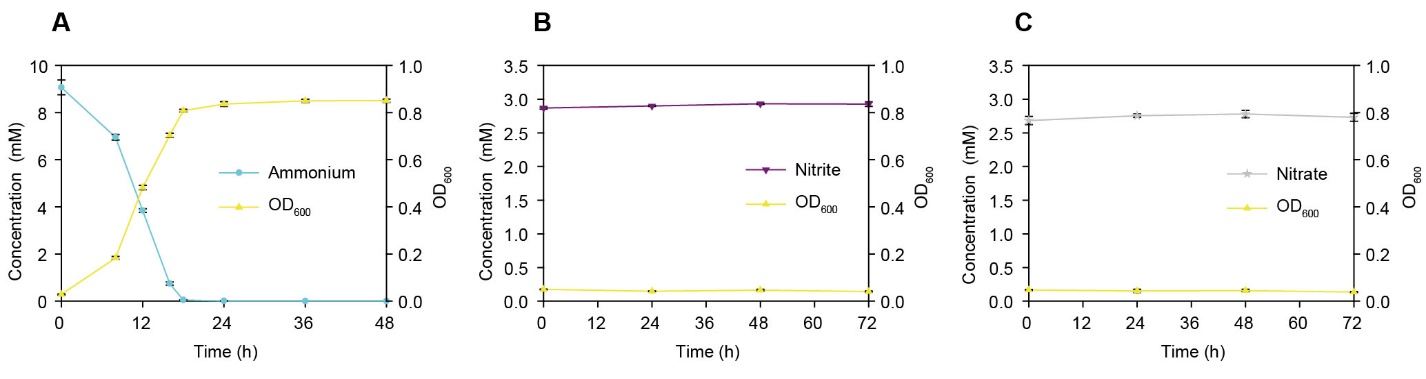

**Fig. S4. Aerobic growth on various nitrogen sources of strain HO-1.** (A) Strain HO-1 grows on ammonia (10 mM) as only nitrogen source. (B) Strain HO-1 does not grow on nitrite (2.8 mM) as nitrogen source. (C) Strain HO-1 does not grow on nitrate (2.8 mM) as nitrogen source. Succinate was used as sole carbon source with a C/N mass ratio of 10 : 1. Values are means ± SD (error bars) for three biological replicates. If not visible, error bars are smaller than symbols.

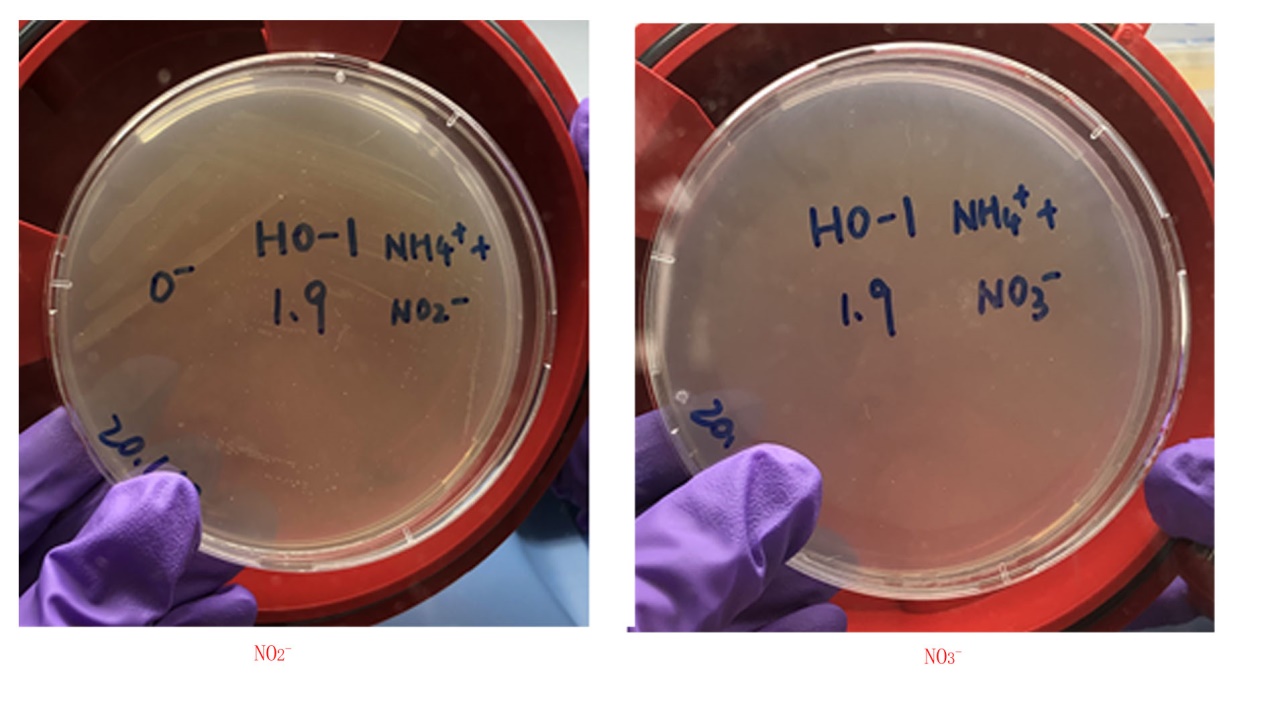

**Fig. S5. Strain HO-1 grew on the HNM medium under anaerobic condition in the presence of ammonium and nitrite (left), but not ammonium + nitrate (right).** HNM media used here included 5mM (^15^NH_4_)_2_SO_4_, 5 mM nitrate or nitrite, and 17.5 mM succinate.

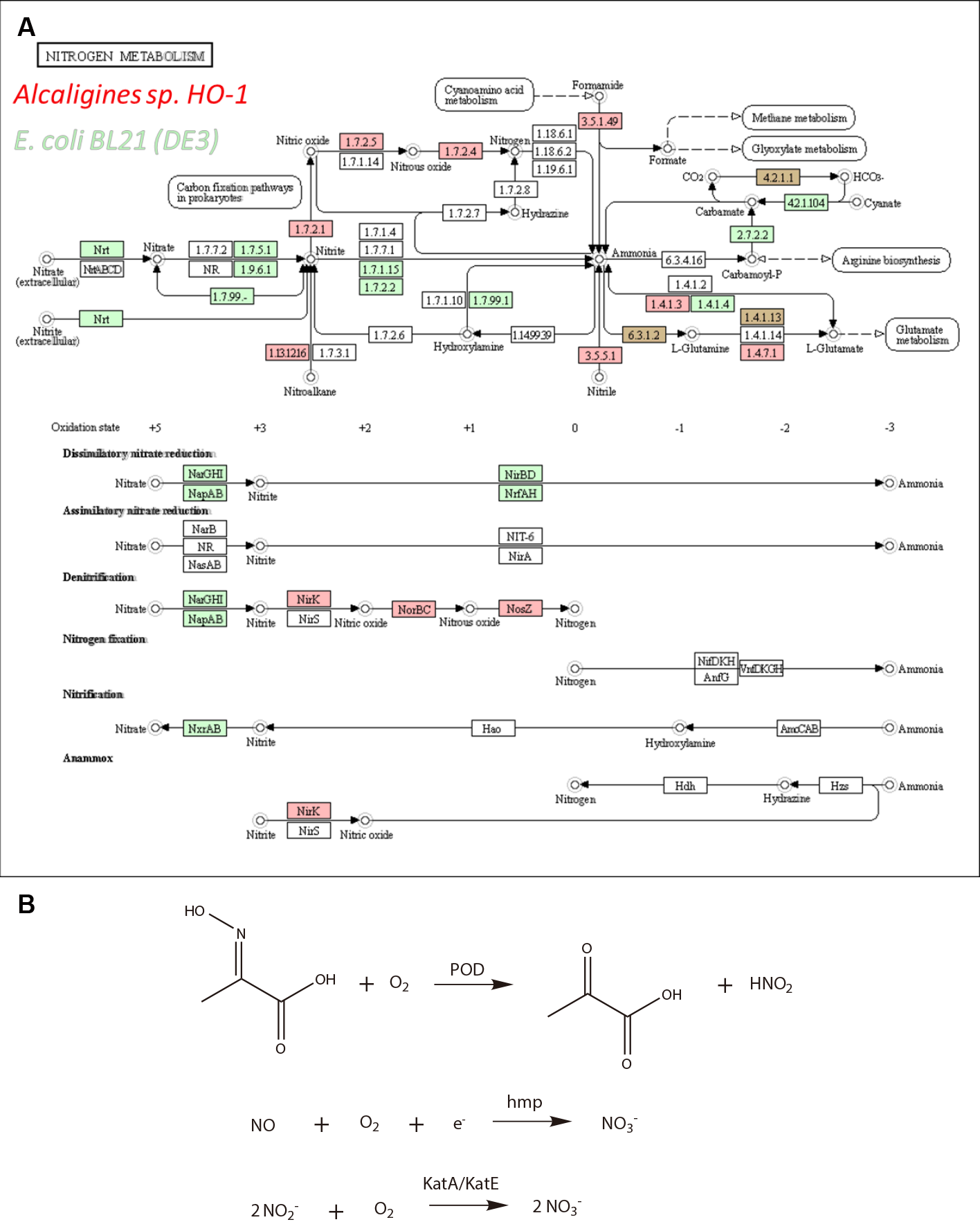

**Fig. S6. Nitrogen metabolism pathway in *Alcaligenes* strain HO-1 and *E. coli* strain BL21.** (a) Predicted nitrogen metabolism according to genome annotation. Homologies in the HO-1 and *E. coli* genome are highlighted in red and green (orange for both). Note that canonical pathways for nitrification, anaerobic ammonium oxidation (anammox), and assimilatory nitrate reduction were missing in strain HO-1. In addition, enzymes for assimilatory or dissimilatory nitrate reduction are absent. The *nxrAB*-like genes in *E. coli* reflect the homology of its *narGH* genes with these genes and do not suggest that *E. coli* oxidizes nitrite. Interestingly, *E. coli* BL21 encodes a hybrid cluster protein that has been associated with hydroxylamine reduction to ammonia (25) and high affinity nitric oxide (NO) reduction to N_2_O (26). (b) Reactions catalyzed by POD, Hmp and KatA/KatE. POD, pyruvic oxime dioxygenase FE795_02445. Hmp, flavohemoglobin-NO-dioxygenase (NO detoxifying enzyme,) FE795_07360 and FE795_16420. KatA/KatE, catalases with nitrite-oxidizing side-activity (KatA FE795_11315 and KatE FE795_17180).

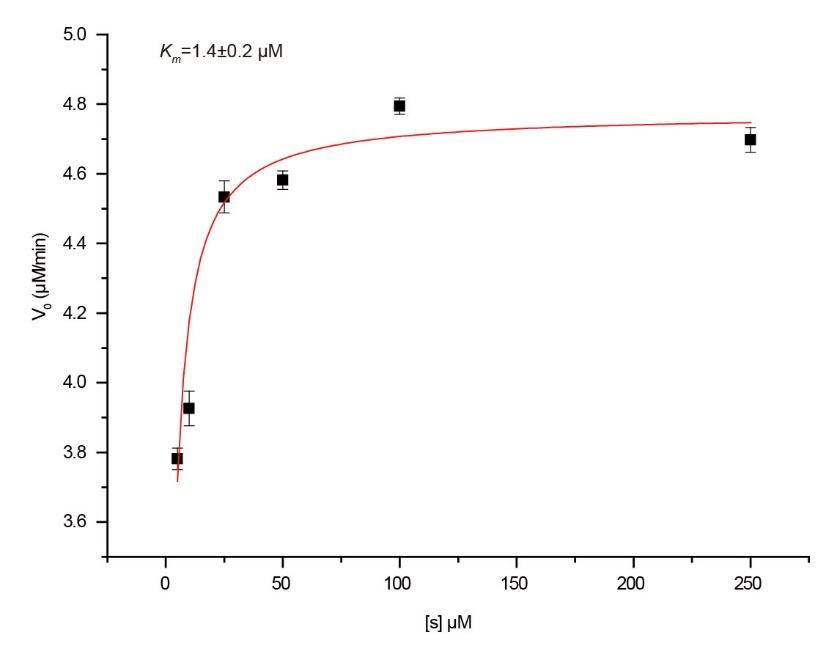

**Fig. S7. Enzymatic assay of NADH oxidation by DnfA in the presence of 5-250 uM hydroxylamine.** The reaction buffer included 10 uM DnfA, 100 uM FAD, 250 uM NADH and 5-250 uM hydroxylamine at atmospheric conditions. The decrease of NADH was monitored by 340 nm on a spectrophotometer at 25 °C. The calculated *K*m is one-order magnitude lower than that determined at 30 °C as described in the main text.

**Dataset S1. (separate file)**

RNA-seq assays of strain HO-1 stimulated by ammonia.
